# Novel Replication Stress-Associated High-Grade Serous Ovarian Cancer Models Recapitulate Diverse Tumour Ecosystems and Enable Preclinical Studies

**DOI:** 10.64898/2026.09.29.755402

**Authors:** V Pourcel, Z Drouin, E Létourneau, Z Gerber, KR Malkoun, D Jean, M Placet, M Labrie

**Author notes:** Corresponding author: Marilyne Labrie, Dept Immunology and Cell Biology, FMSS - Université de Sherbrooke, 3201, Rue Jean-Mignault, Sherbrooke, QC, J1E 4K8.

## Abstract

**Background:** High-grade serous ovarian cancer (HGSC) is characterized by extensive genomic heterogeneity and frequent alterations that promote replication stress (RS). Although RS has emerged as a promising therapeutic vulnerability, it remains unclear whether distinct RS-associated genomic alterations generate similar tumour states or drive unique biological programs and therapeutic dependencies.

**Methods:** We developed and characterized a panel of syngeneic HGSC models representing recurrent RS-associated alterations, including overexpression of *Ccne1*, *Brd4*, *Myc, Ndrg1*, *Pik3ca*, and *Brca1* heterozygosity, alongside previously established *Pten*/*Nf1*-deficient and *Myc* overexpressing models. Tumour fitness, transcriptomic programs, histopathology, extracellular matrix composition, immune organization, and therapeutic vulnerabilities were assessed using subcutaneous (s.c.) and intraperitoneal (i.p.) allografts, RNA sequencing (RNAseq), mass spectrometry (MS), cyclic-immunofluorescence (Cyc-IF), and preclinical therapeutic studies.

**Results:** RS-associated genotypes exhibited marked differences in tumour fitness and generated distinct tumour ecosystem states despite sharing a common genetic background. Tumours harbouring *BRCA1* heterozygosity displayed reduced tumour fitness, mesenchymal features, extracellular matrix remodelling, stromal expansion, and immune-excluded phenotypes. In contrast, tumours co-expressing *Ccne1* and *Brd4* exhibited the highest RS level together with strong activation of DNA damage response pathways and limited immune infiltration. Immune characterization identified immune-desert, immune-suppressed, and immune-excluded states across models, demonstrating substantial genotype-dependent immune heterogeneity. Elevated CD47 expression emerged as a defining feature of the *Ccne1/Brd4* tumour state and was confirmed in human HGSC specimens with high CCNE1 and BRD4 protein expression. Therapeutic targeting of ATR and CD47 in intraperitoneal *Ccne1/Brd4* allografts reduced tumour burden and induced histopathological responses compared with either monotherapy alone, supporting the translational relevance of this RS-high tumour state.

**Conclusions:** Distinct RS-associated alterations do not converge on a common HGSC phenotype but instead generate diverse tumour ecosystem states with unique effects on tumour fitness, stromal remodelling, immune organization, and therapeutic vulnerabilities. These findings highlight the importance of considering the genetic origin of RS when developing therapeutic strategies and establish a framework for investigating genotype-specific vulnerabilities in HGSC.

## Introduction

High-grade serous ovarian carcinoma (HGSC) is the most common and aggressive subtype of epithelial ovarian cancer. HGSC is marked by high heterogeneity, which contributes to its high mortality rate and to the persistent lack of effective, durable treatment options^1–4^. Cytoreductive surgery combined with cytotoxic therapy based on carboplatin and paclitaxel represents the main treatments for HGSC^3,4^. This standard treatment remains highly ineffective, with 80% of patients relapsing within the first three years^3,4^.

Inspired by the paradigm-shifting role of homologous recombination deficiency (HRD) in breast cancer treatment, homologous recombination (HR) status has emerged as an important stratification factor in HGSC. Approximately 50% of HGSC tumours exhibit HRD-related genetic alterations and demonstrate improved initial responses to platinum-based chemotherapy, while some patients also benefit from PARP inhibitors (PARPi)^5^. PARPi exploit synthetic lethality in HRD tumours through the accumulation of unrepaired DNA damage^6,7^. Although these therapies have significantly improved progression-free survival in HRD patients, they provide limited benefit to HR competent (HRC) tumours^5,8^. However, substantial biological heterogeneity persists within both HRD and HRC groups, suggesting that HR status alone does not fully capture the diversity of HGSC biology and therapeutic vulnerabilities.

Beyond HR deficiency, HGSC is characterized by numerous recurrent genomic alterations that can promote replication stress (RS). These include *RB1* and *NF1* loss, amplification of *CCNE1*, *BRD4*, *MYC*, *CDK1*, and *CDK2*, as well as alterations in *PIK3CA* and *CDK12*^2,9^. While RS promotes tumour evolution and genomic diversification, it simultaneously creates a therapeutic vulnerability by increasing reliance on DNA damage response (DDR) pathways and G2/M checkpoints, thereby sensitizing cancer cells to DDR inhibition^10^. Consequently, inhibitors targeting ATR, CHK1, PKMYT1, and WEE1 have been investigated in HGSC^10,11^. Tumours harbouring *CCNE1* amplification, for example, appear particularly sensitive to several G2/M checkpoint inhibitors alone or in combination^11–13^. Importantly, these alterations induce RS through distinct biological mechanisms, and it remains unclear whether they generate equivalent tumour states or drive unique biological programs associated with different therapeutic dependencies. Clinically, major challenges remain in identifying predictive biomarkers for DDR inhibitor response and in selecting patients most likely to benefit from these therapies while limiting toxicity^14^.

In addition to tumour-intrinsic alterations, HGSC is characterized by a complex tumour microenvironment (TME) that contributes to treatment resistance. Only a minority of HGSC tumours are considered immunologically “hot,” displaying robust immune infiltration and anti-tumour response associated with improved clinical outcomes^15–18^. In contrast, many tumours exhibit an immune-excluded phenotype in which stromal components form physical and biochemical barriers that limit immune cell infiltration and function^16,19^. HGSC tumours can also evade anti-tumour immunity through the expression of immune checkpoint molecules such as PD-1, PD-L1 and CTLA-4. However, immune checkpoint blockade has demonstrated only modest efficacy in HGSC, with response rates generally below 15%^20^.

Accumulating evidence suggests that tumour genotype can profoundly influence not only cell-intrinsic oncogenic signalling, but also tumour architecture, epithelial-mesenchymal plasticity, stromal remodelling, immune infiltration, and therapeutic response. However, the extent to which common HGSC driver alterations shape distinct tumour ecosystems remains poorly understood. Dissecting these relationships has been challenging because of the limited availability of genetically controlled and immunocompetent preclinical models that permit direct comparison of individual oncogenic events within a shared genetic background. To address this challenge, we investigated recurrent HGSC alterations, including several associated with RS, using a syngeneic HGSC platform composed of newly engineered and previously validated models. New genotypes were generated to represent *CCNE1*, *BRD4*, *MYC*/*NDRG1* and *PIK3CA* alterations with or without BRCA1 heterozygosity and were analysed alongside the previously described PPNM and BPPNM models^21^ derived from the same parental Trp5^−/−^ lineage^21,22^. Using transcriptomic, histopathological, and spatial proteomic analyses, we sought to determine how specific oncogenic alterations influence tumour fitness, tumour ecosystem organization, and immune composition. Our results demonstrate that distinct HGSC genotypes drive divergent tumour ecosystem states characterized by differences in tumour initiation, epithelial-mesenchymal plasticity, stromal remodelling, immune organization, and therapeutic vulnerabilities.

## MATERIAL AND METHODS

### Clinical Samples

Clinical HGSC specimens were collected in accordance with institutional ethical guidelines and approved research protocols, with informed consent obtained from all participants.

### Generation and culture of HGSC cell lines

The models used in this study deribed from murine fallopian tube epithelial cell lines carrying *Trp53* deletion (*Trp53*^−/−^) or combined *Trp53* deletion and *Brca1* heterozygosity (*Trp53*^−/−^; *Brca1*^+/−^), as previously described^21,22^. To generate additional HGSC models, cells were transduced with lentiviral vectors expressing mutant TRP53*^R172H^*together with combinations of *Ccne1*, *Myc*, *Brd4*, *Pik3ca*, and *Ndrg1*. TRP53*^R172H^*was expressed from a pLV-EF1α-IRES-Hygro vector, whereas all additional cDNAs were cloned into customized pHAGE-PURO (Addgene, #118692) or pHAGE-BLAST lentiviral vectors and sequence-verified prior to use. Lentiviral particles were produced in HEK293T cells using standard packaging and envelope plasmids and used to transduce target cells. Transduced cells were selected using hygromycin, puromycin, and/or blasticidin as appropriate. All cell lines were maintained in DMEM supplemented with HEPES, EGF, and 5% heat-inactivated fetal bovine serum (FBS) and were routinely confirmed to be free of Mycoplasma contamination by PCR.

### Immunofluorescence

For immunofluorescence staining, cells were seeded onto circular glass coverslips placed in 24 well plates and cultured for 24 to 48 h to reach up to 70% confluence. Once the desired confluence was achieved, the culture medium was removed and the coverslips were washed with phosphate-buffered saline (PBS).Cells were fixed with 3% paraformaldehyde (PFA, Sigma Aldrich #P6148) for 15 min at room temperature (RT), followed by permeabilization with 0.1% Triton X-100 in PBS for 5 min at RT. Non-specific binding sites were blocked using 1%bovine serum albumin (BSA) in PBS for 1 h at RT. Primary antibodies diluted in 1%BSA/PBS were incubated either for 2 hours at RT or overnight at 4°C. After washing with PBS, samples were incubated with the appropriate secondary antibodies for 1 hour at RT. Nuclei were stained with Hoechst 33342 (Invitrogen, #R37605) at a final concentration of 10 μg/mL for 5 min at RT. Coverslips were mounted using Immu-Mount mounting medium (epredia #9990402) and allowed to solidify before imaging. Fluorescence images were acquired using an Axioscope 5 fluorescence microscope (Zeiss). A complete list of primary and secondary antibodies used for immunofluorescence experiments is provided in **Supplementary Table 1**.

### Animal Studies

All animal experiments were approved by the Institutional Animal Research Review Committee of the Université de Sherbrooke and conducted in accordance with Canadian Council on Animal Care guidelines. Female C57BL/6 mice (6-8 weeks old; Charles River Laboratories) were used for all studies. For subcutaneous allografts, 1 × 10⁷ cells were injected into both flanks in Matrigel and tumour growth was monitored three times weekly by caliper measurements. Tumour volume was calculated using the formula [(length × width²)/2]. Mice were euthanized upon reaching predefined humane endpoints including tumour ulceration. For intraperitoneal allografts, 1 × 10⁷ cells were injected intraperitoneally in PBS and mice were monitored three times weekly for weight loss and clinical signs of disease. Animals were euthanized when humane endpoints were reached and tumours were collected for downstream characterization.

### Therapeutic studies

For treatment studies, mice bearing established intraperitoneal tumours were randomized to treatment groups four weeks after tumour implantation (n=10 mice per group). Anti-CD47 antibodies (MIAP301, Bioxcell) were administered intraperitoneally three times weekly for six weeks (200µg per injection). Ceralasertib was administered daily by oral gavage at 50 mg/kg using an intermittent schedule (1 week on, 1 week off) for six weeks. Control animals received vehicules alone consisting of rat IgG2a isotype control (clone 2A3, Bioxcell) and propylene glycol (Bioshop). Mice were euthanized one day after the 6 weeks of treatment. Mice weights were acquired throughout the experiment to assess treatment tolerability.

### Tissue processing and tissue microarray construction

Tumour tissues were washed in PBS, fixed overnight in 3.7% formaldehyde, and paraffin embedded. FFPE blocks were sectioned into 4 μm sections by the Histology Facility of the Université de Sherbrooke. For tissue microarray (TMA) construction, 3 to 4 representative cores (1.5 mm diameter) from each tumour (n=3-4 independent tumours per genotype) were transferred into recipient paraffin blocks. TMA sections were subsequently used for multiplex immunofluorescence analyses.

### Cyclic-immunofluorescence

Cyc-IF was performed, by HORBITUS Platform at Université de Sherbrooke, as previously described^23–25^. Briefly, FFPE tissue sections were deparaffinized, rehydrated, and subjected to antigen retrieval using sequential citrate (pH 6.0) and Tris-EDTA (pH 9.0) buffers. Following blocking, tissue autofluorescence was acquired using an Axioscan 7 slide scanner (Zeiss). Sequential cycles of staining, imaging, and fluorophore inactivation were performed using primary antibodies directly conjugated to Alexa Fluor 488, 555, 647, or 750 fluorophores. Fluorescence signal inactivation was achieved using hydrogen peroxide and sodium hydroxide treatment between cycles. Images were acquired after each staining cycles. A complete list of antibodies is provided in **Supplementary Table 2**. Following image acquisition, image registration was performed using ASHLAR^26^. Cell segmentation, feature extraction, and single-cell data export were performed using QI Tissue Image Analysis software (v1.4.0). Quality control included the removal of aberrant cells based on nuclear size and autofluorescence levels. Marker intensities were corrected for autofluorescence at the single-cell level and normalized using z-score transformation. Cell populations were identified based on lineage-marker expression profiles. Spatial coordinates generated during segmentation were used for tissue reconstruction and spatial analyses of tumour and stromal compartments. For spatial compartment analyses, TMA cores were subdivided into 200 × 200-pixel grids and classified as tumour-rich or stromal regions based on cellular composition. Data processing, statistical analyses, and visualization were performed using custom Python scripts or SCORPy^27^.

### RNA sequencing

RNA sequencing was performed on n = 3-4 independent tumours per genotype. Tumour samples preserved in RNAlater were homogenized and total RNA was extracted using the RNeasy Mini Kit (Qiagen) according to the manufacturer’s instructions. RNA quality and concentration were assessed by spectrophotometry prior to library preparation and sequencing at the CHU de Québec-Université Laval Genomics Platform. Libraries were sequenced on a NovaSeq 6000 platform. Sequencing reads were aligned to the mouse reference genome (GRCm39/mm39) using STAR, and transcript abundance was quantified using Salmon. Lowly expressed genes detected in fewer than 10% of samples were excluded from downstream analyses. Gene-level expression matrices were generated by aggregating transcripts sharing the same annotated gene symbol. To estimate pathway activity at the sample level, Gene set variation analysis (GSVA) was performed in R using Hallmark and Reactome gene sets obtained from MSigDB. Pathways represented by fewer than five genes were excluded. Transcriptomic data processing, statistical analyses, and visualizations were performed using custom R and Python scripts.

### Mass spectrometry

Mass spectrometry (MS) was performed on n = 3-4 independent tumours per genotype. Tumour tissues were homogenized and proteins were extracted in urea-based lysis buffer. Following protein quantification, 50 μg of protein per sample were reduced, alkylated, and digested overnight with trypsin. Peptides were purified using C18 solid-phase extraction and analyzed by liquid chromatography coupled to tandem MS (LC-MS/MS) using a nanoElute HPLC system interfaced with a timsTOF Pro mass spectrometer (Bruker Daltonics). Data were acquired in diaPASEF mode. Raw DIA files were processed using DIA-NN (v1.8.1) against the UniProt mouse proteome database (UP000000589). Protein identification was performed using a library-free workflow with *in silico* spectral library generation and deep-learning-based prediction of spectra, retention times, and ion mobility. The resulting Unique.Gene report, provided by the MS-based proteomics Facility of the Université de Sherbrooke, was used for downstream analyses. Proteins detected below an empirically defined detection threshold or in fewer than 10% of samples were excluded. All subsequent data processing, statistical analyses, and visualizations were performed using custom Python 3 scripts implemented in Jupyter Notebook.

### Statistical analysis

Statistical analyses were performed using GraphPad Prism, R, or Python. Comparisons between two groups were performed using two-tailed unpaired Student’s t-tests. Comparisons involving multiple groups were performed using one-way or two-way analysis of variance (ANOVA). Survival analyses were performed using Kaplan-Meier estimates and compared using the log-rank test. Unless otherwise indicated, data are presented as mean ± SEM. Statistical significance was defined as P < 0.05.

## RESULTS

### Establishment of syngeneic models representative of distinct HGSC genotypes associated with RS

HGSC is genetically heterogeneous, although some alterations are known to be frequent and sometimes co-occurring. Analysis of the publicly available Firehose Legacy cohort highlights *TP53* as the major common alteration in HGSC, being altered in 88% of samples (**Fig.1A**). *BRCA1* and *BRCA2*, the principal determinants of HRD, are altered in 8% and 12% of tumours, respectively. Among alterations associated with RS, *CCNE1* and *BRD4* are amplified in 26% and 28% of patients, respectively. Although these genes are located on distinct *loci* on chromosome 19, their amplification significantly co-occurs (**Supplementary Table 3**). Furthermore, *CCNE1* is mutually exclusive with *BRCA* alterations, as previously described^28^. *MYC* is amplified in 43% of cases and significantly co-occurs with *NDRG1* amplification, consistent with their shared chromosomal locus at 8q24 (**Supplementary Table 3**). Finally, *PIK3CA* is amplified in nearly 36% of the samples. These recurrent alterations represent genetically distinct routes to HGSC development and provide an opportunity to investigate how specific oncogenic events influence tumour behaviour and tumour ecosystem organization.

**Figure 1.**
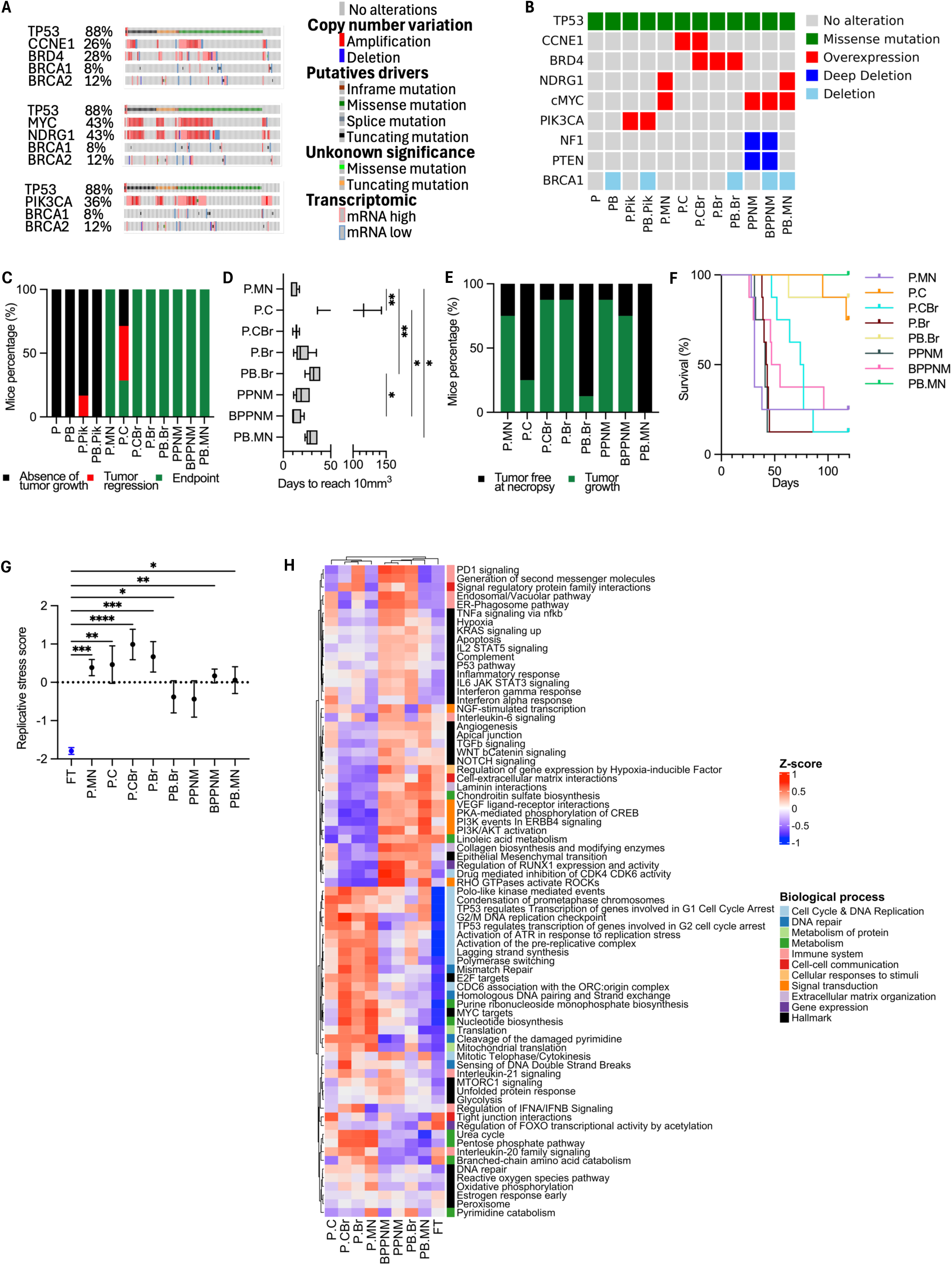
RS-associated HGSC alterations differentially shape tumour fitness and oncogenic programs. **(A)** OncoPrint showing the frequency and co-occurrence of recurrent genetic alterations in human HGSC from the TCGA Firehose Legacy cohort. **(B)** OncoPrint showing the combinations of HGSC-associated genetic alterations represented across the 12 syngeneic murine models used in this study. **(C)** Percentage of successful subcutaneous (s.c.) tumour formation for each allograft model. **(D)** Time required for s.c. tumours to reach 10 mm³ following implantation. **(E)** Percentage of successful intraperitoneal (i.p.) tumour formation for each allograft model. **(F)** Kaplan-Meier survival curves of mice bearing i.p. tumours derived from each model. **(G)** RS score derived from bulk RNA-seq across allograft models and fallopian tube controls (FT). **(H)** Heatmap of GSVA enrichment scores derived from bulk RNA-seq showing the most variable pathways across allograft models based on sorted MSigDB Hallmark (H) and Reactome (C2). Pathways are grouped according to biological function. For panels D and G, data are presented as median ± SEM; *P < 0.05, **P < 0.01, *** P<0,0001, ****P<0,00001 (one-way ANOVA test). n = 3-4 tumours per group for RNA-seq analyses, n = 4-6 tumours per group for s.c. growth analyses, and N = 8 mice per group for i.p. studies.

To model these clinically relevant genomic alterations, we generated and validated derivatives models of the previously described Trp53^−/−^ syngeneic HGSC platform^21,22^ (**Fig.1B**, **Supplementary Fig.1A-F**). All generated cell lines harbour the TRP53^R172H^ mutation, corresponding to the human TP53^R175H^ hotspot mutation frequently observed in HGSC^29,30^. Cell lines overexpressing RS-associated genes *Ccne1* and/or *Brd4* (P.C, P.Br, P.CBr), *Pik3ca* (P.Pik), or *Myc* and *Ndrg1* (P.MN) were generated from the parental *Trp53^−/−^* cell line to model the co-amplification patterns observed in human tumours. For clinically relevant genotypes, additional derivative carrying heterozygous *Brca1* loss (*Trp53^−/−^,Brca^+/−^* parental cell line) were established (PB.Br, PB.MN, and PB.Pik) to investigate the impact of *Brca1* heterozygosity. The previously published BPPNM and PPNM models were also incorporated to expand the spectrum of RS-associated alterations represented in the platform^21^. Together, these twelve cell lines constitute a genetically controlled platform encompassing recurrent HGSC alterations and varying degrees of genomic instability.

### BRCA1 heterozygosity reduces tumour fitness in RS-associated HGSC models

RS can promote tumour evolution through genetic instability, but also impose substantial stress on cancer cells, creating a balance between tumour-promoting and tumour-limiting effects^31,32^. Because recurrent HGSC alterations induce RS through distinct mechanisms, their impact on tumour fitness may differ substantially. In addition, RS-associated alterations frequently exhibit non-random patterns of co-occurrence or mutual exclusivity with BRCA1 alterations, suggesting a complex interplay between RS and HR pathways. Whether BRCA1 heterozygosity influences the fitness of tumours driven by distinct RS-associated oncogenic alterations remains poorly understood. We therefore evaluated the tumorigenic potential of each genotype *in vivo*. Each cell line was injected subcutaneously into immunocompetent C57BL/6 mice, providing a controlled, reproducible model for tumour characterisation (**Fig.1C-D**). The P and PB cell lines (**Fig.1B**) failed to generate tumours, demonstrating that Trp53^R172H^ alone is insufficient to drive tumour formation in either a BRCA1 wild-type or heterozygous context (**Fig.1C**). Overexpression of PIK3CA alone exhibited very limited tumorigenic capacity. No tumours formed in the PB.Pik model, while only 16% of P.Pik allografts initially established before rapidly regressing and being cleared by the host. These findings suggest that *Pik3ca* overexpression alone is insufficient to sustain tumour growth and likely requires additional oncogenic alterations to promote stable tumorigenesis as already describe in different context^33,34^. In contrast, *Brd4* overexpression was sufficient to induce tumour formation in both *Brca1* wild-type and heterozygous backgrounds, with 100% tumour incidence observed in P.Br and PB.Br mice. Interestingly, 42% of P.C tumours regressed following implantation. Moreover, tumour initiation was significantly delayed in the P.C model compared with the other genotypes (**Fig.1D**, **Supplementary Fig.1G**). These observations are consistent with the notion that excessive *Ccne1*-driven RS imposes a fitness cost that can limit tumour establishment^35^. In contrast, the P.CBr model exhibited 100% tumour formation and substantially shorter latency, suggesting that *Brd4* overexpression mitigates the detrimental consequences associated with *Ccne1*-induced RS (**Fig.1C-D**, **Supplementary Fig.1G**).

Comparison of matched genotypes further revealed that tumour fitness was strongly influenced by *Brca1* status. In both the *Brd4* and *Myc*-*Ndrg1* backgrounds, *Brca1* heterozygosity delayed tumour initiation relative to the corresponding *Brca1* wild-type models. Tumour onset occurred significantly earlier in P.MN compared with PB.MN, and a similar trend was observed when comparing P.Br and PB.Br (**Fig.1D**, **Supplementary Fig.1G**). Notably, these differences in latency were not accompanied by major changes in subsequent tumour growth kinetics, indicating that *Brca1* heterozygosity predominantly affects tumour establishment rather than tumour expansion (**Supplementary Fig.1G**). In contrast, no obvious effect of *Brca1* heterozygosity was observed in the PPNM/BPPNM pair in the subcutaneous setting (**Fig.1D**), suggesting that the consequences of partial *Brca1* loss depend on the accompanying oncogenic landscape.

The effects observed in the subcutaneous model were consistently more pronounced in the intraperitoneal allograft model (**Fig.1E,F**). The toxicity associated with *Ccne1* overexpression was further accentuated, with only 25% of P.C allografts resulting in significant slower tumour progression and significantly prolonged survival compared with P.Br and P.CBr (**Fig.1F, Supplementary Fig.2A)**. As observed subcutaneously, co-expression with *Brd4* mitigated this phenotype, increasing the tumour uptake rate of P.CBr allografts to 87.5%, comparable to that observed in the P.Br model. P.Br-bearing mice displayed shorter survival than P.CBr-bearing mice, notably associated with the development of ascites in more than 60% of the animals (**Fig.1E,F, Supplementary Fig.2A**). The impact of *Brca1* heterozygosity was particularly evident in the intraperitoneal setting. While P.Br and P.MN readily established intraperitoneal tumours, fewer than 12.5% of PB.Br and PB.MN allografts resulted in tumour growth, leading to significant prolonged survival (**Fig.1E,F, Supplementary Fig.2A**). Furthermore, intraperitoneal tumours derived from P.Br and P.MN displayed significantly higher tumour progression scores compared to their *Brca1* heterozygous-deleted counterparts, PB.Br and PB.MN, respectively, although the difference in tumour weight was significant only for the P.Br/PB.Br pair (**Supplementary Fig.2B-D)**. These findings indicate that *Brca1* heterozygosity reduces tumour fitness across multiple RS-associated HGSC genotypes. Interestingly, this phenotype was not observed in the *Pten*/*Nf1*-deficient background. More than 75% of both PPNM and BPPNM allografts generated intraperitoneal tumours (**Fig.1E**), which were associated with ascites formation in over 85% mice bearing tumours. The accumulation of ascitic fluid frequently led to earlier humane endpoint criteria being reached, thereby shortening survival in these cohorts (**Fig.1F, Supplementary Fig.2A**). Together, these observations suggest that the effect of BRCA1 heterozygosity on tumour fitness is highly context dependent and can be overridden by additional tumour-promoting alterations.

**Figure 2.**
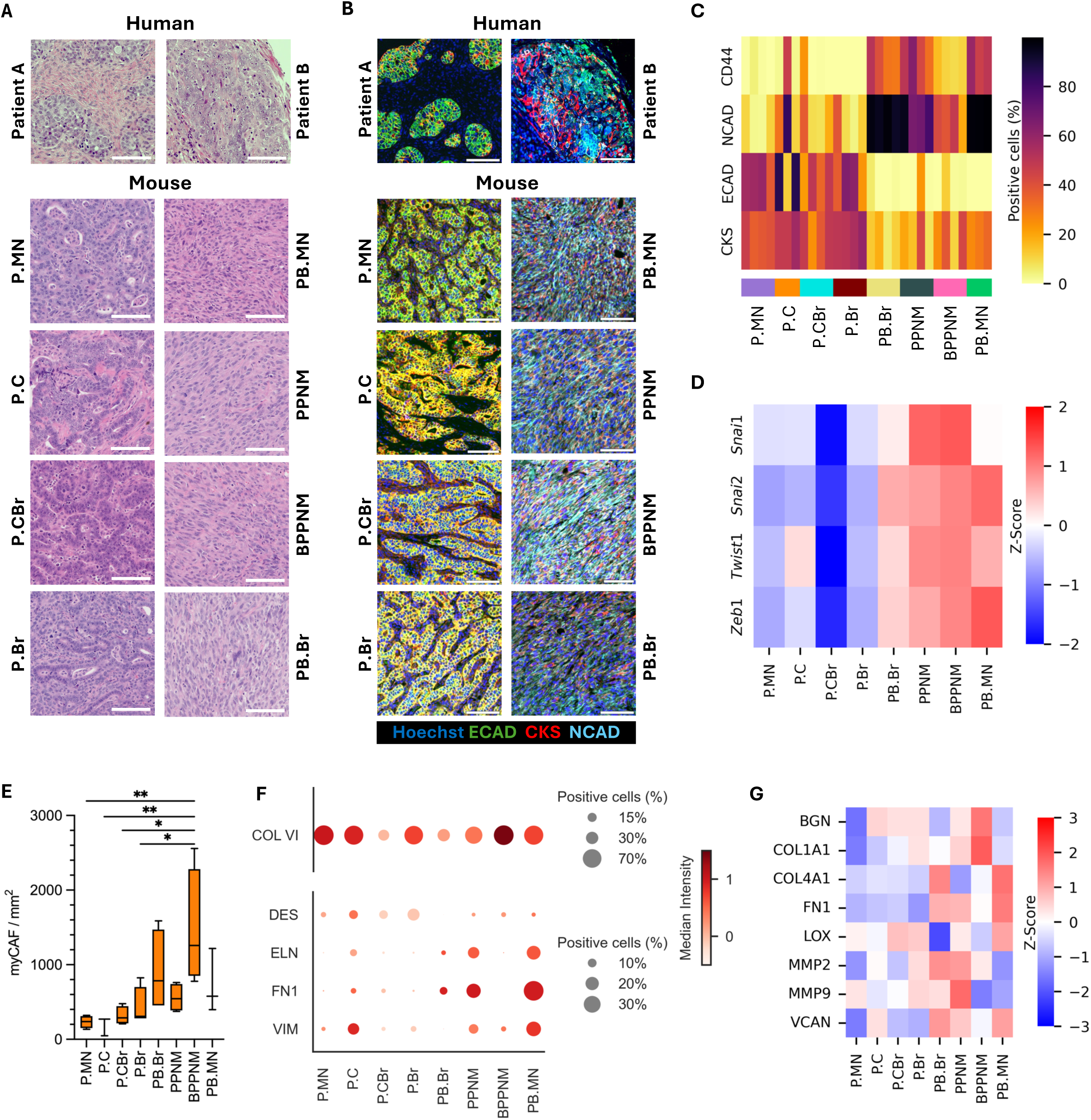
Tumour genotype shapes histological features and ECM composition. **(A)** Representative hematoxylin and eosin (H&E)-stained sections of human HGSC specimens and syngeneic allograft models. Scale bars, 100 μm. **(B)** Representative Cyc-IF images showing E-cadherin (green), cytokeratin 8/19 (red), and N-cadherin (cyan) expression in human HGSC specimens and murine allografts. Nuclei were counterstained with Hoechst (blue). Images were acquired from regions corresponding to the H&E sections shown in panel A. Scale bars, 100 μm. **(C)** Heatmap showing the percentage of cancer cells expressing epithelial (CKs, ECAD) and mesenchymal (CD44, NCAD) markers as determined by Cyc-IF analysis. **(D)** Heatmap showing relative expression of EMT-associated transcription factors measured by bulk RNA-seq. **(E)** Cancer-associated myofibroblast (myCAF) densities across allograft models determined by Cyc-IF analysis. **(F)** Dot plot showing the percentage of stromal cells expressing stromal-associated markers and the corresponding median marker intensity determined by Cyc-IF analysis. **(G)** Heatmap showing the relative abundance of ECM proteins measured by MS. Data in panel E are presented as median ± SEM; *P < 0.05, **P < 0.01 (one-way ANOVA test). n = 3-4 tumours per group for Cyc-IF, RNAseq and MS analyses.

### RS-associated genotypes define distinct transcriptional states

Given the marked differences in tumour fitness observed across genotypes, we next sought to determine whether these alterations generated distinct levels of RS and induced specific transcriptional programs. To enable a comprehensive comparison of genotype-specific tumour states, subsequent molecular analyses were performed on subcutaneous allografts. Unlike the intraperitoneal model, where several genotypes exhibited limited tumour take rates, the subcutaneous setting provided sufficient tumour material across the different genetic backgrounds while preserving interactions with the host microenvironment. This approach enabled direct comparison of the biological consequences of distinct RS-associated alterations independently of differences in tumour dissemination and engraftment efficiency.

To quantify RS, we performed RNA sequencing (RNA-seq) on each tumors and applied the RNA-based RS signature developed by Takahashi et al.^36^ to all subcutaneous allografts and used fallopian tube tissue as a control (**Fig.1G**). Interestingly, RS levels were significantly higher in all allografts than in fallopian tube tissue. Among all genotypes, P.CBr displayed the highest RS score. In contrast, tumours harbouring *Brca1* heterozygosity and/or *Pten* loss exhibited slightly lower RS scores than the remaining allografts. These findings indicate that although all engineered alterations generate RS-associated tumours, the magnitude of RS differs depending on the underlying oncogenic driver. To further characterize the biological consequences of these alterations, we performed gene set variation analysis (GSVA) on RNA-seq data using fallopian tube tissue as a control (**Fig.1H**). Visualization of the analysis as a clustered heatmap revealed three major groups. Fallopian tube tissue formed a distinct cluster separate from all tumour samples. The remaining allografts segregated into two major tumour clusters. One cluster grouped tumours harbouring *Brca1* heterozygosity and/or *Pten* loss (BPPNM, PPNM, PB.Br, and PB.MN), while the second cluster contained P.C, P.CBr, P.Br and P.MN tumours. This segregation was notable because tumours clustered according to broader biological programs rather than individual engineered alterations, suggesting the emergence of distinct transcriptional states. Within the *Brca1* heterozygous/*Pten*-deficient cluster, pathways associated with epithelial-mesenchymal transition (EMT) and extracellular matrix (ECM) remodelling were strongly enriched, including apical junction, epithelial-mesenchymal transition, laminin interactions, collagen biosynthesis and collagen modifying enzymes. Comparison of P.Br with PB.Br and P.MN with PB.MN further revealed that *Brca1* heterozygosity was associated with reduced enrichment of oxidative stress-related pathways, including reactive oxygen species, oxidative phosphorylation, peroxisome and pentose phosphate metabolism, as well as decreased activity of several metabolic programs involved in amino acid catabolism, nucleotide biosynthesis and the urea cycle. Conversely, pathways involved in cell-cycle progression, DNA replication and DNA repair were broadly enriched across all allografts relative to fallopian tube tissue. Notably, the G2/M DNA replication checkpoint, ATR activation in response to RS and DNA double-strand break sensing pathways were most strongly represented in tumours with wild-type *Brca1* and *Pten*, with the strongest enrichment observed in the P.CBr model. Together, these analyses reveal that recurrent HGSC alterations generate distinct transcriptional states characterized by different levels of RS and activation of divergent biological programs.

### *Brca1* heterozygosity and *Pten* loss define a mesenchymal tumour state

The transcriptional analyses identified a cluster of tumours characterized by *Brca1* heterozygosity and/or *Pten* loss that exhibited strong enrichment of EMT- and ECM-related pathways. To determine whether this transcriptional state was reflected at the histological and protein levels, we performed a comprehensive characterization of tumour architecture and cellular phenotype. Histopathological analysis of all allografts revealed that the models recapitulate the major architectural patterns observed in human HGSC, including solid, papillary, cribriform and tubular structures (**Fig.2A**, **Supplementary Table 4**). The allografts displayed predominantly epithelioid and fusiform cytology, supporting their representativeness of human HGSC^37^. Despite their common HGSC features, the tumours segregated into two major histopathological groups. Tumours harbouring *Brca1* heterozygosity and/or *Pten* loss displayed a predominantly tubular architecture, whereas the remaining genotypes more frequently exhibited papillary and cribriform structures. Importantly, these phenotypes are consistent with those observed in human HGSC (**Fig.2A)** and these histological patterns were largely conserved in the intraperitoneal model (**Supplementary Fig.2E**).

Because tubular architecture has been associated with epithelial-mesenchymal transition in other cancers^38^ we next examined the expression of epithelial and mesenchymal markers using cyclic-immunofluorescence (Cyc-IF). All tumours were stained for the epithelial markers cytokeratin 8/19 (CKS) and E-cadherin (ECAD), together with the mesenchymal markers N-cadherin (NCAD) and CD44 (**Fig.2B**). Quantification of marker expression in cancer cells (**Fig.2C**) demonstrated that tumours harbouring *Brca1* heterozygosity and/or *Pten* loss expressed higher levels of NCAD and CD44, whereas the remaining allografts displayed increased expression of CKS and ECAD. These findings confirm that the histological differences observed between tumour clusters are associated with distinct epithelial and mesenchymal cellular states. This observation is consistent with previous reports identifying BRCA1 and PTEN as negative regulators of EMT^39–41^.

To determine whether these phenotypic differences resulted from transcriptional reprogramming, we examined the expression of EMT-associated transcription factors using RNA-seq data (**Fig.2D**). EMT-related transcription factors were more highly expressed in tumours harbouring *Brca1* heterozygosity and *Pten* loss, a trend that was particularly apparent when comparing PB.Br with P.Br and PB.MN with P.MN. Conversely, P.CBr tumours exhibited lower expression of EMT-associated transcription factors than either P.C or P.Br tumours, suggesting that combined *Ccne1* and *Brd4* overexpression promotes maintenance of a more epithelial state. Importantly, these results validate the transcriptional clustering identified in **Fig.1H** and suggest that specific RS-associated alterations promote different tumour identities.

### Mesenchymal tumour states exhibit enhanced stromal remodelling and ECM deposition

Because EMT is frequently associated with stromal activation and ECM remodelling, we next investigated whether the mesenchymal tumour state identified above was accompanied by broader ecosystem changes. Analysis of the tumour cellular composition revealed modest differences in the density of major cell populations across allografts (**Supplementary Fig.3**). *Brca1* heterozygous deleted and *Pten* deficient allograft tended to exhibit lower tumour cell density (**Supplementary Fig.3A**), reaching significance between PB.NM and P.MN, P.CBr, P.Br and between PB.Br and P.CBr. This trend was complemented with increased stromal cell densities (**Supplementary Fig.3B**) in *Brca1* heterozygous and *Pten* deficient allograft. The stromal enrichment was characterized by the presence of cancer-associated myofibroblasts (myCAFs), which was significantly higher in BPPNM compared to allografts without *Pten* or *Brca1* alterations (**Fig.2E**). Comparison of P.Br and PB.Br as well as P.MN and PB.MN further suggested that *Brca1* heterozygosity contributes to stromal expansion (**Supplementary Fig.3B**). Endothelial cell densities remained similar across genotypes (**Supplementary Fig.3C**). similar levels of vascularization. Likewise, immune cell densities were largely comparable across models (**Supplementary Fig.3D**), with the exception of a significant difference between BPPNM and PB.MN tumours, identifying these genotypes as the most immune-inflamed and immune-desert phenotypes, respectively. However, the functional significance of these differences could not be inferred from total immune cell density alone and was therefore investigated in subsequent analyses.

**Figure 3.**
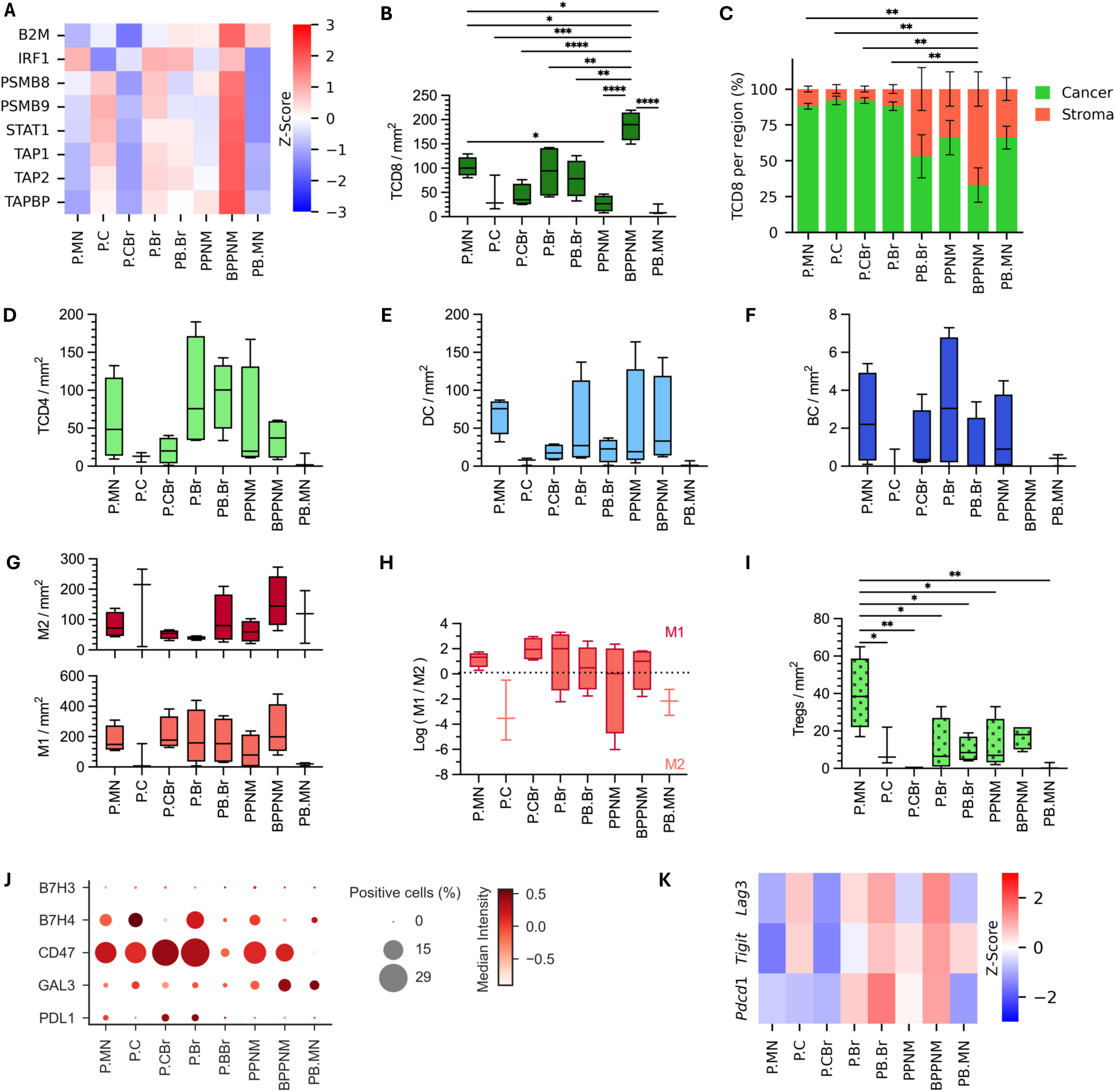
RS-associated genotypes shape distinct immune ecosystems and immune regulatory programs. **(A)** Heatmap showing the relative abundance of proteins involved in antigen presentation measured by MS. **(B)** CD8⁺ T-cell densities across allograft models determined by single-cell Cyc-IF analysis. **(C)** Distribution of CD8⁺ T cells within tumour and stromal compartments determined by grid-based spatial analysis of Cyc-IF data. **(D-G)** Immune cell densities (cell/mm^2^) of **(D)** CD4⁺ T cells, **(E)** dendritic cells, **(F)** B cells and **(G)** M1-like and M2-like macrophages across allograft models as determined by single-cell Cyc-IF analysis. **(H)** M1/M2 macrophage ratio across allograft models. **(I)** Regulatory T-cell (Treg) densities across allograft models determined by single-cell Cyc-IF analysis. **(J)** Dot plot showing the percentage of cancer cells expressing immunoregulatory molecules and the corresponding median marker intensity determined by Cyc-IF. **(K)** Heatmap showing relative expression of T-cell exhaustion markers measured by bulk RNA-seq. Data are presented as median ± SEM. *P < 0.05, **P < 0.01, *** P<0,0001, ****P<0,00001 (one-way ANOVA). n = 3-4 tumours per group for Cyc-IF, RNAseq and MS analyses.

Given the differences observed in overall stromal cell density across genotype, we next characterized the composition of the ECM using Cyc-IF. Collagen VI was the most abundant stromal component across all allografts and was particularly enriched in BPPNM tumours (**Fig.2F**). In contrast, desmin and elastin were expressed at comparatively lower levels across all genotypes. Expression of fibronectin and vimentin varied more substantially between tumour models, with elevated levels observed in P.C, PPNM and PB.MN tumours. To determine whether these protein-level observations reflected a broader remodelling of the ECM, we analysed the expression of additional ECM-associated proteins using mass spectrometry (MS) (**Fig.2G**). Tumours harbouring *Brca1* heterozygosity and/or *Pten* loss expressed a more diverse repertoire of ECM components and generally displayed higher expression levels than the remaining genotypes. This distinction was particularly evident when comparing P.MN and PB.MN tumours, with PB.MN exhibiting substantially greater expression of ECM-associated proteins such as metalloproteinases (MMPs), collagen and fibronectin. In contrast, P.MN tumours displayed relatively low and poorly diversified ECM proteins. The strong ECM signature observed in PB.MN may in part reflect lower expression of matrix metalloproteinases MMP2 and MMP9, limiting matrix turnover and remodelling. Together, these findings demonstrate that the mesenchymal tumour state identified in *Brca1*-heterozygous and *Pten*-deficient tumours extends beyond cancer cell-intrinsic EMT programs and is accompanied by substantial stromal remodelling. Increased myCAF abundance, enhanced ECM diversity and elevated matrix-associated transcriptional programs collectively support the existence of a distinct stromal-rich ecosystem state associated with these genotypes.

### RS-associated genotypes shape distinct immune ecosystems

Because immune infiltration and activation are strongly influenced by both tumour-intrinsic programs and the surrounding stromal environment, we next investigated whether the distinct transcriptional and stromal states identified above were associated with differences in immune ecosystem composition and organization. Analysis of proteins involved in antigen presentation by MS revealed substantial differences across tumour genotypes (**Fig.3A**). Among all models, BPPNM tumours displayed the strongest activation of antigen presentation programs, with robust expression of multiple proteins involved in MHC-I processing and antigen presentation. In contrast, weaker antigen presentation signatures were observed in P.MN, PB.MN and P.CBr tumours. Interestingly, PB.Br tumours exhibited reduced expression of PSMB8 and PSMB9 despite maintaining expression of the remaining antigen presentation-associated proteins, suggesting incomplete activation of the immunoproteasome pathway. To determine whether these differences were associated with altered immune infiltration, we quantified CD8⁺ T-cell densities across all models (**Fig.3B**). BPPNM tumours exhibited a significant higher density of CD8⁺ T cells compared with all other models consistent with BPPNM being immune-inflamed. All other allograft displayed low level of CD8+ T cells infiltration with PB.MN displayed the lowest level, consistent with an immune-desert phenotype. Interestingly, although CD8^+^ T cell abundance was high in BPPNM, we examined their localization within tumour and stromal compartments (**Fig.3C**) and found that CD8⁺ T cells were predominantly localized within stromal regions, suggesting a physical barrier restricting access to tumour nests. A higher stromal localization was also observed in the other models harbouring *Brca1* heterozygosity or *Pten* loss, however only BPPNM reached statistical significance compared to remains genotypes. These findings suggest that stromal remodelling induce by those alterations may influence immune access to cancer cells.

Analysis of additional immune populations revealed further differences between tumour genotypes. CD4⁺ T-cell densities varied considerably across models, ranging from nearly undetectable in P.BMN tumors levels to a median of approximately 100 cells/mm² in PB.Br (**Fig.3D**). Dendritic cells were also heterogeneously distributed, with densities ranging from 0 in PB.MN to a median of approximately 75 cells/mm² in P.MN (**Fig.3E**). These observations indicate that antigen-presenting cell recruitment differs substantially between RS-associated genotypes and may contribute to the distinct immune states observed across models. Interestingly, B cells were scarce in all tumour models (**Fig.3F**), indicating that lymphoid infiltrates were predominantly composed of T-cell populations rather than B cells. As macrophages represented a substantial proportion of the immune infiltrate, we next examined macrophage polarization. Quantification of M1-like and M2-like macrophages revealed marked differences between tumour genotypes (**Fig.3G**). Except for P.C and PB.MN which had almost no detectable M1-like macrophages, all models displayed high M1-like densities with an average of approximately 200 cells/mm^2^. Interestingly, the M1/M2 ratio (**Fig.3H**), revealed that P.C and PB.MN were enriched in M2-Like macrophages, consistent with an immunosuppressive pro-tumoral ecosystem.

To further explore immune evasion mechanisms, we calculated the density of Tregs across the models and found that P.MN microenvironment was strongly enriched in Tregs with a median density of 38 cells/mm^2^, suggesting an additional layer of immune suppression despite the presence of infiltrating lymphocytes (**Fig.3I**). We next investigated the expression of clinically relevant immune checkpoint molecules by Cyc-IF (**Fig.3J**). PD-L1 expression was uniformly weak across all tumour models, with only a minimal proportion of tumour cells expressing PD-L1. The highest average expression was observed in PCBr and PBr tumours, where PD-L1-positive cells represented less than 2.5% of the tumour population. B7H3 expression was also weak across models with less than 1% of tumour cells being positive for its expression. In contrast, B7H4 was more broadly expressed, with elevated levels observed in P.MN, P.C, P.Br and PPNM tumours, in which B7-H4-positive cells represented approximately 5% of the tumour cell population on average. Galectin-3 (gal-3) has recently gained attention in HGSC as an immunoregulatory molecule that contributes to an immunosuppressive tumour microenvironment and may represent a novel therapeutic target^42^. Although gal-3-positive cancer cells consistently represented less than 10% of the tumour population, gal-3 expression varied across allograft models, highlighting heterogeneity among RS-associated genotypes. Beyond lymphocyte-associated immune checkpoints, we also assessed the macrophage checkpoint protein CD47, which is known for inhibiting macrophage-mediated phagocytosis. CD47 is frequently overexpressed in HGSC and has been associated with poor clinical outcomes, making it an attractive therapeutic target^43^. CD47 was highly expressed across most genotypes, suggesting a potentially important role for macrophage-dependent immune evasion in these models. Notably, P.MN, P.C, P.CBr, P.Br, PB.Br and PPNM tumours displayed the highest levels of CD47, with up to 30% of cancer cells being positive for the protein. These findings indicate that distinct RS-associated genotypes preferentially engage different immune regulatory pathways and highlight several genotype-specific therapeutic opportunities. To further investigate immune dysfunction, we assessed the expression of immune exhaustion-associated genes using RNA-seq (**Fig.3K**). *Pdcd1*, *Tigit* and *Lag3* expression was elevated in P.Br, PB.Br and BPPNM tumours, suggesting the presence of exhausted T-cell populations. Together, these observations indicate that the nature of immune regulation differs substantially between RS-associated genotypes.

To integrate the immune parameters evaluated across the model panel, including antigen presentation, immune cell composition, spatial organization and immune regulatory programs, we classified tumours into distinct immune ecosystem states (**Table 1**). P.C, P.CBr and PB.MN tumours were characterized by low antigen presentation, limited CD8⁺ T-cell infiltration, scarce dendritic cells and immunosuppressive myeloid features, consistent with an immune-desert phenotype. In contrast, P.Br and P.MN tumours displayed greater lymphocyte infiltration but were associated with evidence of immune regulation, including elevated Treg abundance in P.MN and increased expression of immune checkpoint and exhaustion-associated markers in P.Br, consistent with an immune-suppressed state. Finally, PB.Br, PPNM and BPPNM tumours exhibited stromal-rich microenvironments associated with retention of CD8⁺ T cells within stromal compartments despite detectable immune infiltration, indicative of an immune-excluded phenotype. Together, these findings demonstrate that RS-associated oncogenic alterations do not generate a uniform immune landscape but instead establish distinct immune ecosystem states that may require fundamentally different therapeutic strategies.

**Table 1.** Immune features associated to each syngeneic model genotype.

| **Models** | **Immune features** | **Proposed phenotypes** |
| --- | --- | --- |
| **P.MN** | Low antigen presentation<br>High Tregs abundance | **Immune suppressed** |
| **P.C** | Limited CD8+ T-cells infiltration<br>Low dendritic cells infiltration<br>Immunosuppressive myeloid features | **Immune desert** |
| **P.CBr** | Low antigen presentation<br>Limited CD8+ T-cells infiltration<br>Low dendritic cells infiltration | **Immune desert** |
| **P.Br** | Increased expression of immune checkpoint<br>Increased exhaustion-associated markers | **Immune suppressed** |
| **PB.Br** | Low dendritic cells infiltration<br>CD8+ T cells stromal retention<br>ECM enriched | **Immune-excluded** |
| **PPNM** | Limited CD8+ T-cells infiltration<br>ECM enriched | **Immune-excluded** |
| **BPPNM** | High antigen presentation<br>CD8+ T cells stromal retention<br>Increased exhaustion-associated markers<br>CAF-Enriched<br>ECM enriched | **Immune-excluded** |
| **PB.MN** | Low antigen presentation<br>Limited CD8+ T-cells infiltration<br>Low dendritic cells infiltration<br>Immunosuppressive myeloid features<br>ECM enriched | **Immune desert** |

### The high-RS CCNE1/BRD4 tumour state reveals a clinically relevant therapeutic opportunity

The characterization of our RS-associated syngeneic panel demonstrated that distinct genomic alterations generate divergent tumour ecosystem states characterized by differences in tumour fitness, stromal remodelling, immune organization and RS levels. Beyond providing a framework to study HGSC heterogeneity, this platform offers an opportunity to identify genotype-specific therapeutic vulnerabilities and evaluate rational treatment strategies. Among the analysed genotypes, P.CBr emerged as a particularly relevant model. The combined overexpression of *Ccne1* and *Brd4* generated the highest RS levels observed across the panel and recapitulates a clinically important HGSC subgroup associated with poor prognosis and therapeutic resistance^44^. Given their elevated RS, these tumours are expected to exhibit increased dependence on DNA damage response pathways, supporting the rationale for targeting G2/M checkpoint regulators. Although several DDR inhibitors, including ATR inhibitors, have demonstrated promising activity in RS-high tumours, clinical and preclinical studies suggest that combination strategies will likely be required to maximize therapeutic efficacy^45,46^. Current combination approaches involving DDR-targeting agents frequently aim to either further increase genomic damage or potentiate anti-tumour immune responses. During the immune ecosystem characterization of our model panel, CD47 emerged as one of the most highly expressed immunoregulatory molecules in P.CBr tumours ((**Fig.3J**). Indeed, P.CBr subcutaneous allografts displayed significantly higher CD47 expression in cancer cells than all other genotypes (**Fig.4A**). Importantly, this phenotype was maintained in both subcutaneous and intraperitoneal tumours (**Fig.4B**), indicating that elevated CD47 expression represents a stable feature of the P.CBr tumour state.

**Figure 4.**
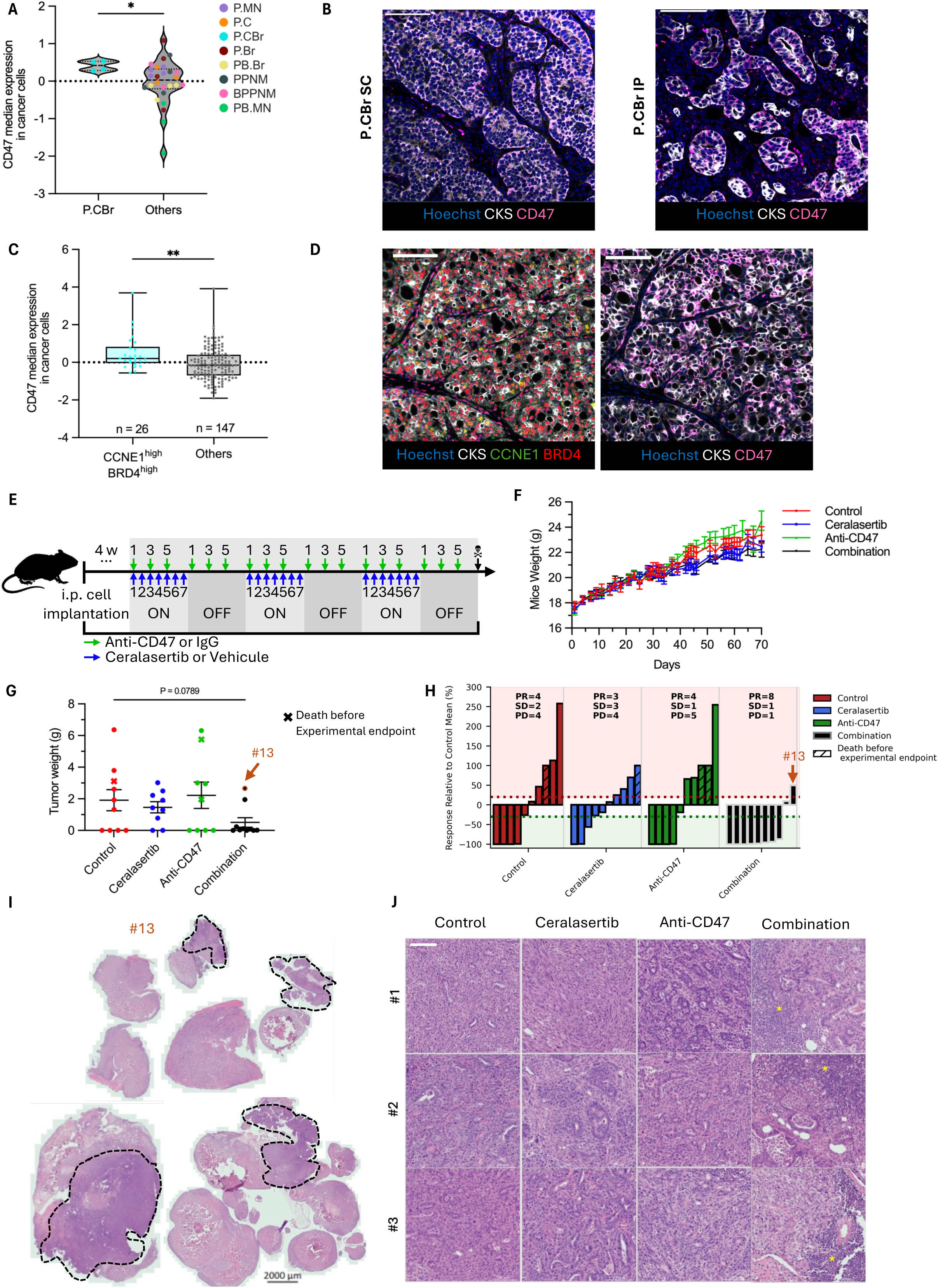
CD47 represents a therapeutic vulnerability in RS-high PCBr tumours. **(A)** CD47 expression in cancer cells from PCBr s.c. allografts compared with all other genotypes, as determined by Cyc-IF analysis. **(B)** Representative immunofluorescence images showing cytokeratin 8/19 (white) and CD47 (pink) expression in s.c. and i.p. PCBr allografts. Nuclei were counterstained with Hoechst (blue). Scale bars, 100 μm. **(C)** CD47 expression in cancer cells from human HGSC specimens with high CCNE1 and BRD4 protein expression compared with all other tumours, as determined by single-cell Cyc-IF analysis. **(D)** Representative Cyc-IF images of a human HGSC specimen with high CCNE1 and BRD4 expression showing CCNE1 (green), BRD4 (red), CD47 (pink), and cytokeratin 8/19 (white). Nuclei were counterstained with Hoechst (blue). Images were acquired from corresponding regions on consecutive tissue sections. Scale bars, 100 μm. **(E)** Experimental design of the therapeutic study. Mice bearing established i.p. PCBr tumours were treated with anti-CD47 or IgG, with ceralasertib or vehicule, or with the combination for six weeks before endpoint assessment. **(F)** Body weight monitoring throughout treatment for each experimental group, n = 10 mice per group (Two-Way ANOVA). **(G)** Tumour weight at experimental endpoint for each treatment group. Crosses indicate mice that reached humane endpoints before study completion. Orange-circled data points correspond to mice #13. **(H)** Waterfall plot showing tumour weight change relative to the mean tumour weight of the control group. Tumours were classified according to RECIST criteria as PR (partial response; −30% change), SD (stable disease: between 20% and −30% change), or PD (progressive disease; +20% change). **(I)** Representative hematoxylin and eosin (H&E)-stained section from mouse #13 treated with the combination therapy. Regions containing viable tumour cells are outlined with a black dashed line. **(J)** Representative H&E-stained sections from three mice per treatment group. Regions of cellular infiltration are indicated by asterisks (*). Scale bars, 100 μm. Data in panel A and C are presented as median ± SEM; *P < 0.05, **P < 0.01 (Mann-Whitney test), (A) n = 3-4 tumours per group, (C) N= 163.

Because CD47 constitutes a major myeloid checkpoint that inhibits macrophage-mediated tumour clearance^43,47^, we next investigated whether this feature was conserved in human disease. Tumours exhibiting co-overexpression of cyclin E1 and Brd4 proteins showed significantly higher CD47 expression than the remaining HGSC cases (**Fig.4C-D**), directly recapitulating the observations made in the syngeneic models. This subgroup accounted for 14.9% of the cohort and was associated with shorter progression-free survival (**Supplementary Fig.4**), consistent with the poor clinical outcome reported for *CCNE1*/*BRD4*-amplified tumours^44^. Together, these findings identify elevated CD47 expression as a clinically relevant hallmark of the high-RS *CCNE1*/*BRD4* tumour state and provide a rationale for combining DDR-targeting agents with CD47 blockade.

### Combining ceralasertib and anti-CD47 in P.CBr allografts induces pathological responses and reduces tumour burden

To validate the therapeutic vulnerabilities identified in our model panel and assess their translational relevance, we performed a preclinical therapeutic study in mice bearing i.p. P.CBr allografts. RS was targeted using the ATR inhibitor ceralasertib, the most clinically advanced ATR inhibitor currently under evaluation^48,49^, while immune evasion was targeted using an anti-CD47 antibody based on the high CD47 expression observed in P.CBr tumours (**Fig.4A-D**). Mice were treated according to the regimen shown in (**Fig.4E**) and described in the method section. All treatment regimens were well tolerated, with no significant differences in body weight observed throughout the study (**Fig.4F**). At endpoint, mice receiving combination therapy exhibited substantially lower tumour burden than control animals (mean ± SEM: 0.508 ± 0.296 g versus 1.780 ± 0.776 g), corresponding to a 71.5% reduction in mean tumour weight (**Fig.4G**). Median tumour burden was reduced from 1.32 g in controls to 0.055 g in combination-treated animals (95.8% reduction). Although the difference did not reach statistical significance (Mann-Whitney p = 0.0789), tumour weights were consistently shifted toward lower values in the combination group. Ceralasertib and anti-CD47 monotherapies produced more modest effects (1.464 ± 0.365 g and 2.222 ± 0.858 g, respectively), indicating that monotherapies are insufficient to elicit a tumour response (**Fig.4G**).

Despite a tumour take rate of approximately 90% in previous experiments, four control mice exhibited either no detectable tumour or only minimal residual disease during necropsy at experimental endpoint. Histological examination further revealed evidence of spontaneous immune infiltration in a subset of control tumours, suggesting that the rat IgG2a isotype control used in this study may have exerted biological activity in this model. To better capture treatment responses at the individual-animal level, tumour burden was normalized to the mean tumour weight of the control group and visualized as a waterfall plot (**Fig.4H**). This analysis stratified the cohort into three groups based on tumour response relative to baseline: responders (PR, including complete responders), mice with stable disease (SD), and mice with progressive disease (PD). Using this classification, we found that most mice receiving the combination therapy experienced either tumour regression or disease stabilization, whereas responses were less frequent in the monotherapy groups. Overall, eight of ten mice treated with the combination displayed reduced tumour burden relative to the control average, compared with only four mice across the monotherapy-treated cohorts, supporting superior activity of the dual treatment strategy. Interestingly, tumour weight alone underestimated response in certain animals. Notably, mouse #13 (combination group), which appeared as an outlier based on tumour mass measurements, harboured very limited viable tumour tissue despite a relatively large residual lesion (**Fig.4I**). Most of the lesion consisted of non-viable tissue associated with extensive inflammatory infiltration, consistent with a late but biologically meaningful therapeutic response. Accordingly, while tumour weight classified this mouse as progressive disease, histopathological assessment suggested a marked treatment effect. Finally, histopathological evaluation further strengthened evidence of therapeutic efficacy in the combination group. Examination of tumours from combination-treated mice demonstrated substantial immune cell infiltration that was largely absent from untreated tumours (**Fig.4J**). Taken together, these findings indicate that combined ATR and CD47 inhibition is more effective than either monotherapy in the high-replication-stress P.CBr model. Although spontaneous responses in the control group likely reduced statistical power, both the distribution of individual responses and the histopathological analyses support clinically relevant anti-tumour activity of the combination regimen and provide proof-of-concept for co-targeting RS and macrophage immune checkpoints in *CCNE1*/*BRD4*-driven HGSC.

## Discussion

HGSC exhibits substantial genomic heterogeneity, yet many recurrent alterations converge on the induction of RS, creating growing interest in RS as both a biological driver and therapeutic vulnerability. While alterations in genes such as *CCNE1*, *BRD4*, *MYC*, *PTEN*, and *BRCA1* have all been associated with RS^50–55^, it remains unclear whether they converge on common tumour states or if they give rise to distinct biological programs with unique therapeutic vulnerabilities. To address this question, we developed and comprehensively characterized a panel of syngeneic HGSC models representing recurrent RS-associated genotypes within a shared genetic background. Our findings demonstrate that RS-associated alterations do not converge on a common tumour phenotype. Rather, they generate distinct tumour ecosystem states characterized by differences in tumour fitness, epithelial-mesenchymal plasticity, ECM remodelling, immune organization, and therapeutic vulnerability. We further identify *Brca1* heterozygosity as a context-dependent determinant of tumour fitness and tumour state, while revealing that tumours co-expressing *Ccne1* and *Brd4* represent a highly RS-enriched subgroup associated with elevated CD47 protein expression and potential sensitivity to combined ATR and CD47 targeting. Together, these results highlight the importance of considering the genetic origin of RS when studying HGSC biology and developing genotype-informed therapeutic strategies.

A central finding of this study is that RS-associated alterations do not produce a uniform tumour phenotype despite converging on a common biological process. RS is frequently regarded as a shared consequence of diverse oncogenic events and is increasingly used to stratify tumours for therapies targeting the DDR, such as G2/M checkpoint inhibitors^56^. However, emerging evidence suggests that RS can also influence tumour ecosystem composition through effects on immune signalling, inflammation, and tumour-microenvironment interactions, indicating that its consequences extend beyond tumour cell-intrinsic vulnerabilities^57^. While, our analyses revealed considerable heterogeneity across RS-associated genotypes, extending well beyond differences in RS magnitude. While P.CBr tumours exhibited the highest RS levels together with strong activation of DNA replication and DDR pathways, tumours harbouring *Brca1* heterozygosity and/or *Pten* loss displayed a distinct transcriptional program characterized by EMT, ECM remodelling, and stromal expansion. These differences were further reflected at the histopathological level and were accompanied by striking variation in immune composition and spatial organization. Notably, our data support the existence of multiple immune ecosystem states, including immune-desert, immune-suppressed, and immune-excluded phenotypes, despite all models arising from a common cell-of-origin and genetic background. Together, these findings suggest that the biological consequences of RS are highly dependent on the underlying oncogenic alterations that generate it. Although these alterations can converge on common RS state, they induce RS through distinct molecular mechanisms. *CCNE1* amplification promotes unscheduled origin firing and excessive DNA replication^58,59^, whereas *MYC* amplification increases transcriptional and replicative activity^60,61^. In contrast, *BRCA1* and *PTEN* alterations can impair distinct mechanisms involved in genome maintenance and DNA damage responses^62–65^. Such mechanistic differences may explain why RS does not translate into a uniform tumour ecosystem state, despite representing a shared biological phenotype. These alterations not only affect the magnitude of replication stress but also the downstream biological programs that shape tumour architecture, stromal organisation, and immune composition. Rather than defining a unique tumour state, RS appears to represent a broader biological framework within which distinct tumour ecosystems can emerge. This observation may have important implications for patient stratification, as tumours exhibiting comparable levels of RS could nevertheless differ substantially in their microenvironmental organization and, consequently therapeutic vulnerabilities. This is particularly relevant given the growing interest in combining RS-targeted therapies with immunomodulatory approaches, whose efficacy may also depend on the ecosystem state in which RS arises.

Another important finding of this study is the context-dependent impact of *BRCA1* heterozygosity on tumour fitness and tumour ecosystem composition. While *BRCA1* deficiency is traditionally viewed as a tumour-promoting event through its effects on HR and genomic instability^66,67^, many evidence suggests that excessive genomic instability may also impose a fitness cost on cancer cells^68^. Consistent with this model, *Brca1* heterozygosity delayed tumour establishment in multiple RS-associated backgrounds, particularly in the intraperitoneal setting, indicating that partial loss of *Brca1* can compromise tumour fitness under specific oncogenic contexts. Interestingly, this phenotype was not observed in the BPPNM model, despite the presence of *Brca1* heterozygosity. This observation suggests that the consequences of impaired HR are strongly influenced by the accompanying oncogenic landscape and may be mitigated by additional tumour-promoting alterations. The absence of a similar phenotype in the BPPNM model raises the possibility that the *Pten/Nf1*-deficient background engages compensatory mechanisms that mitigate the fitness cost associated with reduced BRCA1 function, although the underlying mechanisms remain to be determined. Beyond its effects on tumour fitness, *Brca1* heterozygosity was consistently associated with the emergence of a mesenchymal and stromal-rich tumour state. Tumours harbouring *Brca1* heterozygosity and/or *Pten* loss displayed increased expression of EMT-associated transcription factors, enrichment of ECM programs, greater stromal abundance, and architectural features characteristic of a mesenchymal phenotype. These observations are consistent with previous studies implicating both BRCA1 and PTEN in the regulation of epithelial plasticity and suggest that alterations in genome maintenance pathways can exert broad effects on tumour ecosystem organization. Together, our findings indicate that the impact of *Brca1* heterozygosity extends beyond DNA repair defects and may contribute to the establishment of distinct tumour states characterized by altered stromal interactions, ECM remodelling, and immune organization.

The ecosystem differences observed across genotypes were particularly evident at the immune level. HGSC is frequently viewed as an immunologically “cold” tumour, which has contributed to the limited success of immune checkpoint inhibitors in unselected patient populations^15–20^. However, our data suggest that this characterization may oversimplify the immune heterogeneity present within HGSC. Despite sharing a common cell of origin and being generated in a similar genetic background, our models developed markedly different immune states, ranging from immune-desert and immune-suppressed ecosystems to highly infiltrated yet immune-excluded tumours. Notably, several of the mesenchymal and stromal-rich models displayed retention of CD8+ T cells within stromal compartments rather than tumour nests, suggesting that physical exclusion of immune cells may represent an important mechanism of immune evasion in these tumours. This observation is consistent with growing evidence that ECM remodelling and stromal expansion can restrict T-cell trafficking and impair anti-tumour immunity^69,70^. Importantly, our findings indicate that immune cell abundance alone may not adequately capture the immune status of HGSC tumours, as the spatial organization of immune populations varied substantially between genotypes. These results support the concept that genotype contributes not only to tumour-intrinsic biology but also to the establishment of distinct immune ecosystems, which may ultimately influence responsiveness to immunotherapy and other immune-modulating approaches.

Among the tumour states identified in our model panel, the high-RS *Ccne1*/*Brd4* genotype emerged as particularly relevant from a translational perspective. Amplification of *CCNE1* defines a clinically important subgroup of HGSC characterized by poor prognosis, primary treatment resistance, and limited therapeutic options^28,71^. Unlike *BRCA1/2*-deficient tumours, most *CCNE1*-amplified tumours are HR proficient and derive limited benefit from chemotherapy and PARP inhibition, creating an important unmet clinical need^71–74^. Consistent with the known role of Cyclin E1 in driving unscheduled DNA replication^28,56^, P.CBr tumours displayed the highest RS levels and strongest activation of DDR pathways across our model panel. In addition, we identified elevated CD47 expression as a defining feature of this tumour state and confirmed this association in human HGSC specimens co-expressing CCNE1 and BRD4. These findings suggest that the biological consequences of RS extend beyond tumour cell-intrinsic vulnerabilities and may also influence mechanisms of immune evasion. Although the therapeutic study did not reach formal statistical significance, likely owing in part to the limited cohort size and unexpected activity in the isotype control group, several observations support the therapeutic potential of combining ATR inhibition with CD47 blockade. Importantly, the antitumour activity observed in the isotype control group represents a limitation of the study, as isotype antibodies are not always biologically inert and can exert Fc-dependent immunomodulatory effects^75,76^. This may have contributed to the spontaneous tumour regressions observed in some control animals, increasing variability and reducing the apparent differences between treatment groups. Combination-treated mice exhibited the lowest tumour burden, the highest proportion of disease stabilization or regression, and histopathological evidence of therapeutic activity that was not fully captured by tumour weight measurements alone. Notably, one apparent outlier retained a large residual mass despite having very limited viable tumour tissue, illustrating the limitations of endpoint tumour weight as the sole measure of response in this setting. Collectively, these findings provide preliminary evidence supporting further investigation of therapeutic strategies combining CD47-targeted approaches with inhibition of the replication stress response in CCNE1/BRD4-driven HGSC. Future studies should evaluate whether similar effects can be achieved using additional inhibitors targeting the G2/M checkpoint and related DNA damage response pathways, including WEE1, PKMYT1, or CHK1 inhibitors. Likewise, assessment of alternative CD47-targeting agents currently under clinical development will be important to confirm the robustness and translational relevance of this therapeutic concept. Validation in additional preclinical models and host backgrounds will further clarify the potential of combining replication stress-targeted therapies with macrophage checkpoint blockade in this clinically challenging subgroup.

## Conclusion

In conclusion, this study demonstrates that RS-associated alterations do not converge on a single HGSC phenotype but instead generate diverse tumour ecosystem states with distinct effects on tumour fitness, epithelial plasticity, stromal remodelling, immune organization, and therapeutic vulnerability. These findings challenge the notion that RS alone is sufficient to define tumour behaviour and highlight the importance of considering the genetic mechanisms underlying its induction. Our results further suggest that *BRCA1* heterozygosity can influence both tumour fitness and ecosystem composition in a context-dependent manner, while identifying the high-RS *CCNE1/BRD4* state as a clinically relevant subgroup with unique biological and therapeutic characteristics. Collectively, our work provides new insight into how recurrent HGSC driver alterations shape tumour evolution and microenvironmental organization and establishes a framework for investigating genotype-specific vulnerabilities. As therapeutic strategies targeting replication stress continue to advance in the clinic, a deeper understanding of the ecosystem consequences of distinct RS-associated genotypes may prove essential for patient stratification and the development of more effective combination therapies.

## Acknowledgments

The authors acknowledge the Animal Care Facility, Mass Spectrometry-Based Proteomics Facility and the histology and spatial biology sections of the HORBITUS Platform at the Université de Sherbrooke as well as the Genomics Platform at CHU de Québec-Université Laval for their technical support and expertise. We also thank Dr. Gordon Mills (Oregon Health & Science University, OR, USA) and his team for their assistance in generating the plasmids used in this study.

## Supplementary Figure Legends

**Supplementary Figure 1.**
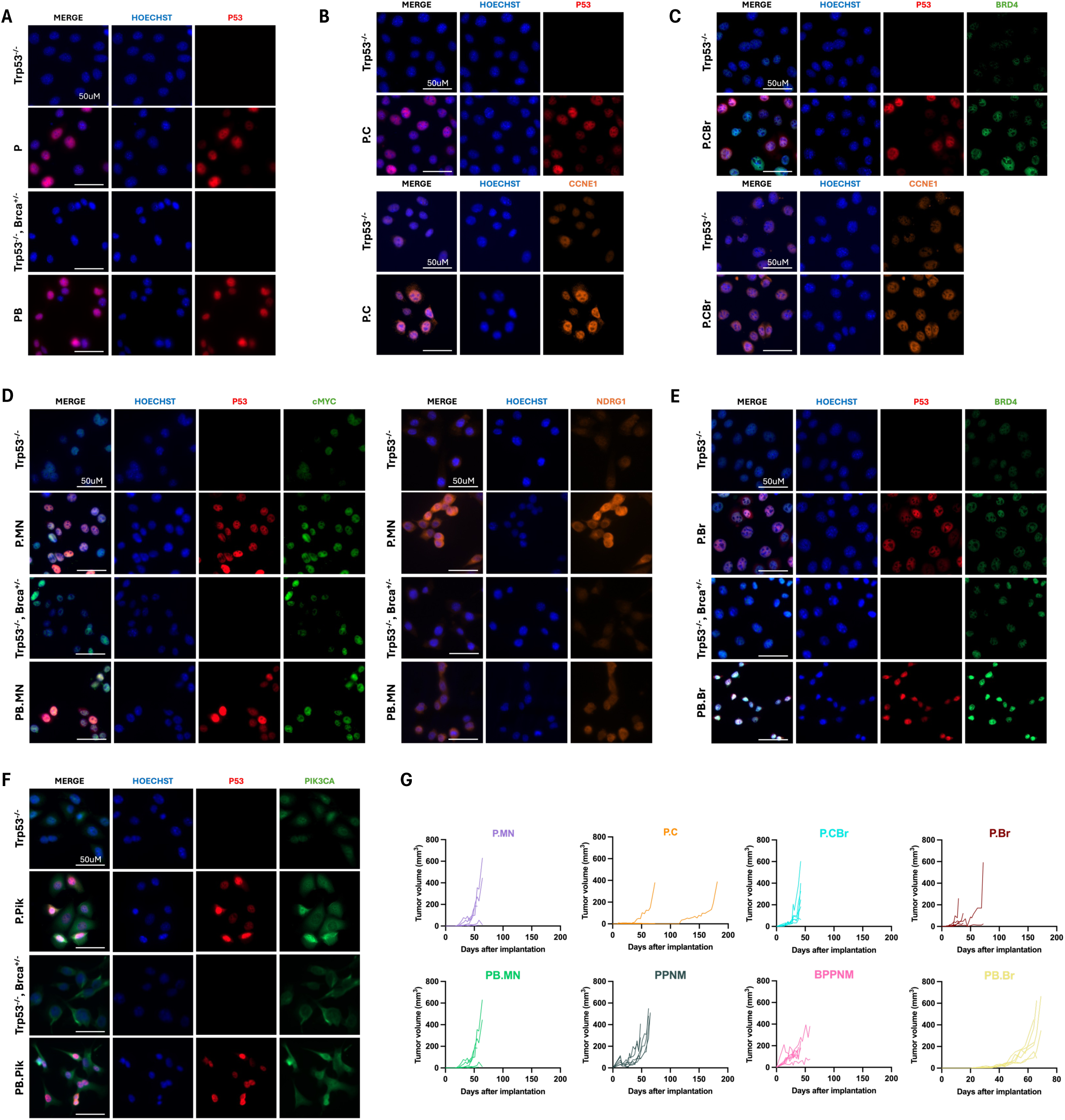
Validation of engineered HGSC cell lines and establishment of s.c syngeneic tumour models. **(A-F)** Immunofluorescence validation of engineered HGSC cell lines. Representative images showing expression of proteins corresponding to the introduced genetic alterations in cell lines **(A)** P and PB cell lines, **(B)** P.C, **(C)** P.CBr, **(D)** P.MN and PB.MN, **(E)** P.Br and PB.Br, and **(F)** P.Pik and PB.Pik. Parental *Trp53^−/−^* and *Trp53^−/−^;Brca1^+/−^* cells were used as controls. Nuclei were counterstained with Hoechst (blue). Scale bars, 50 μm. **(G)** Individual growth curves of successfully established s.c. allografts. Tumour volume was calculated as [(length × width²)/2].

**Supplementary Figure 2.**
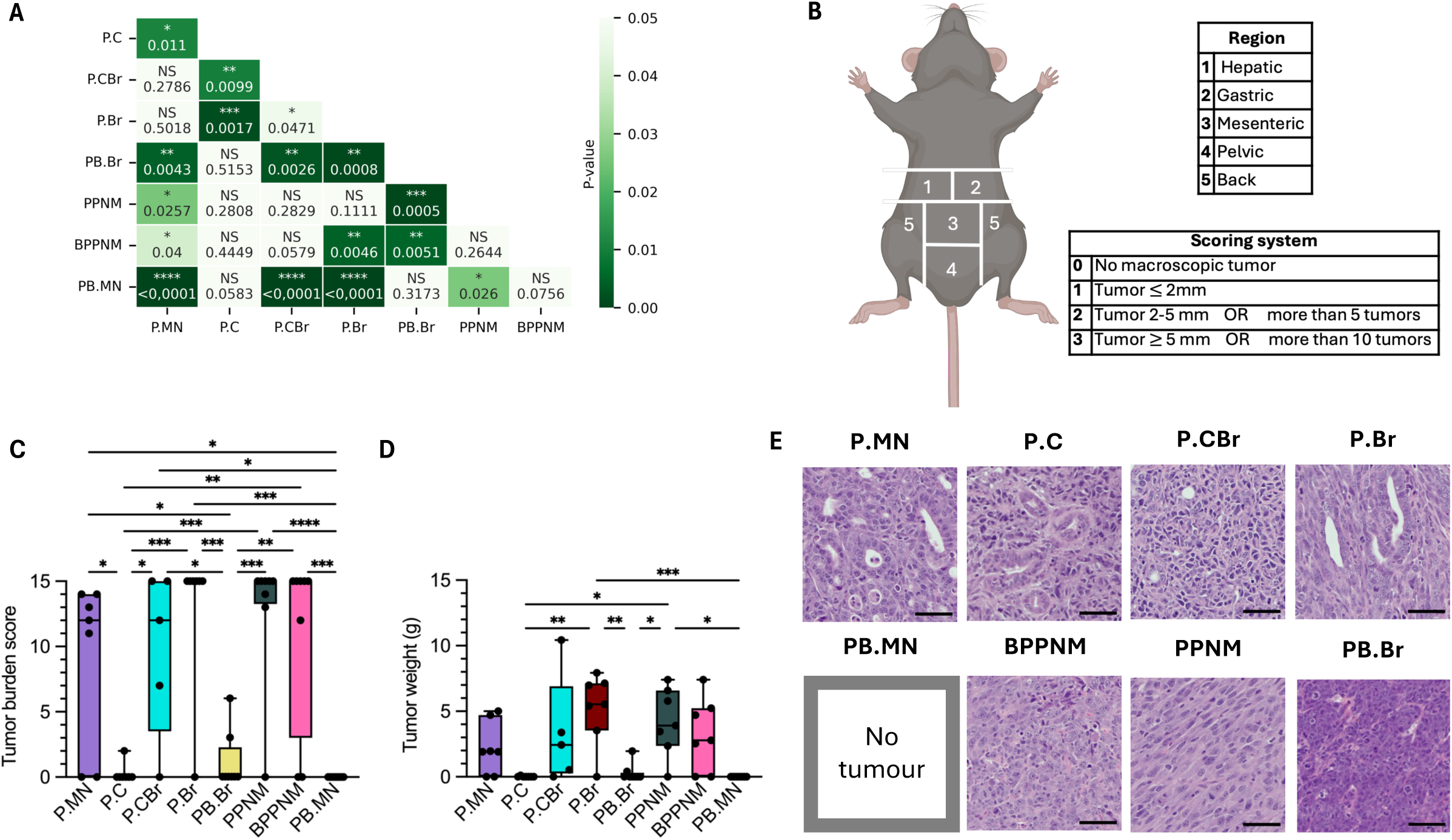
Establishment of i.p syngeneic tumour models. **(A)** Correlation heatmap of statistical significance from Kaplan–Meier survival analysis shown in Fig.1F. Each cell represents the P-Value obtained from the corresponding log-rank test comparing the indicated groups. *P < 0.05, **P < 0.01, ***P < 0.001, and ****P < 0.0001, NS not significant. **(B)** Scoring system inspired by Bastiaenen et al^77^ Peritoneal tumour burden is assessed by dividing the murine peritoneal cavity into five regions and a score equivalent to the amount or the size of tumours per region. Final score correspond to the sum of all regions with a maximum score of 15. **(C)** Tumour burden score across allograft models at humane endpoint. **(D)** Tumour weight across allograft models at necropsies. **(E)** Representative H&E-stained sections of i.p. tumours arising from the indicated allograft models. Scale bars, 50 μm. Data in panel C and D are presented as median ± SEM; *P < 0.05, **P < 0.01, *** P<0,0001, ****P<0,00001 (one-way ANOVA), N = 8 mice per group.

**Supplementary Figure 3.**
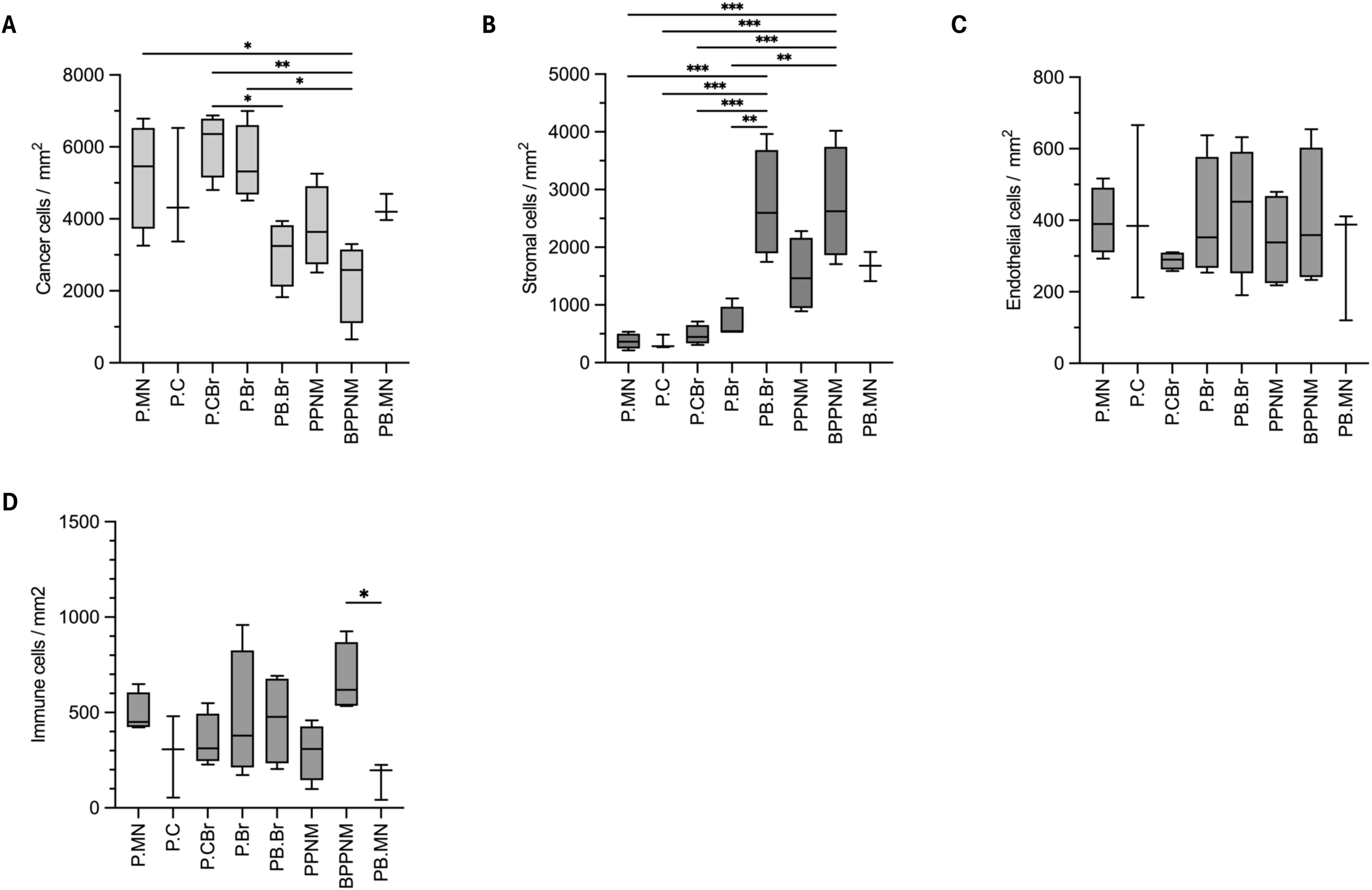
RS-associated genotypes exhibit distinct tumour microenvironment compositions. **(A-D)** Cell population densities (cells/mm^2^) across allograft models determined by single cell Cyc-IF analysis. **(A)** Cancer cells density. **(B)** Stromal cells density. **(C)** Endothelial cells density. **(D)** Total immune cells density. Data are presented as median ± SEM. *P < 0.05 **P < 0.01, *** P<0,0001 (one-way ANOVA), n = 3-4 tumours per group.

**Supplementary Figure 4.**
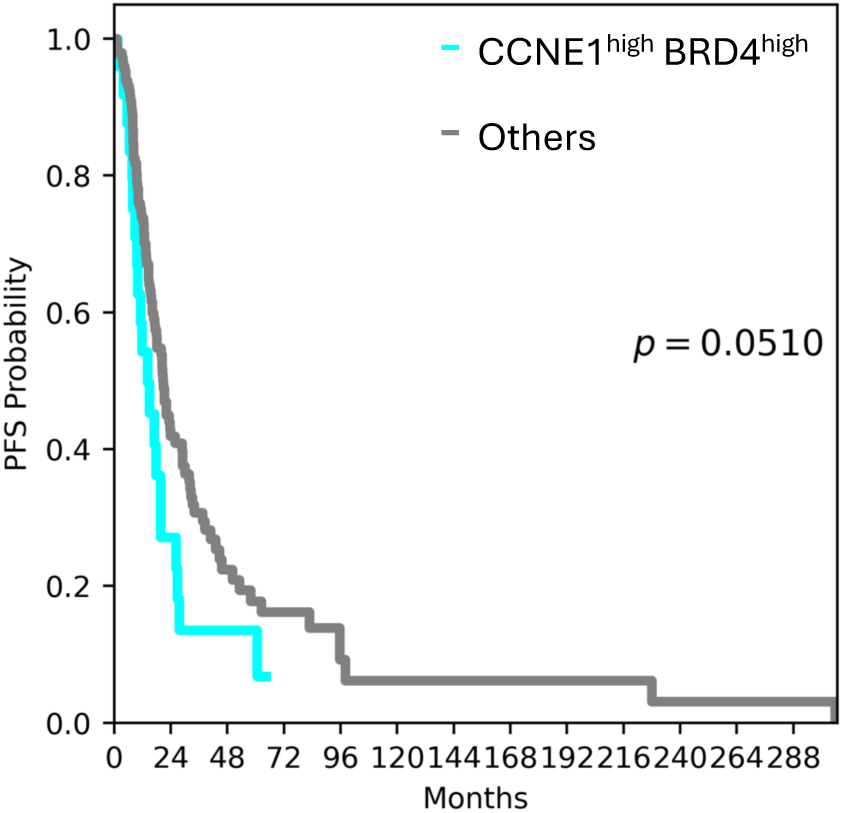
High CCNE1 and BRD4 expression is associated with poorer clinical outcome in human HGSC. Kaplan-Meier analysis of overall survival in patients with HGSC stratified according to CCNE1 and BRD4 protein expression profiles determined by Cyc-IF. Patients with tumours exhibiting high CCNE1 and BRD4 expression were compared with all other patients. Survival distributions were compared using the log-rank test.

---

**Supplementary Table 1.**
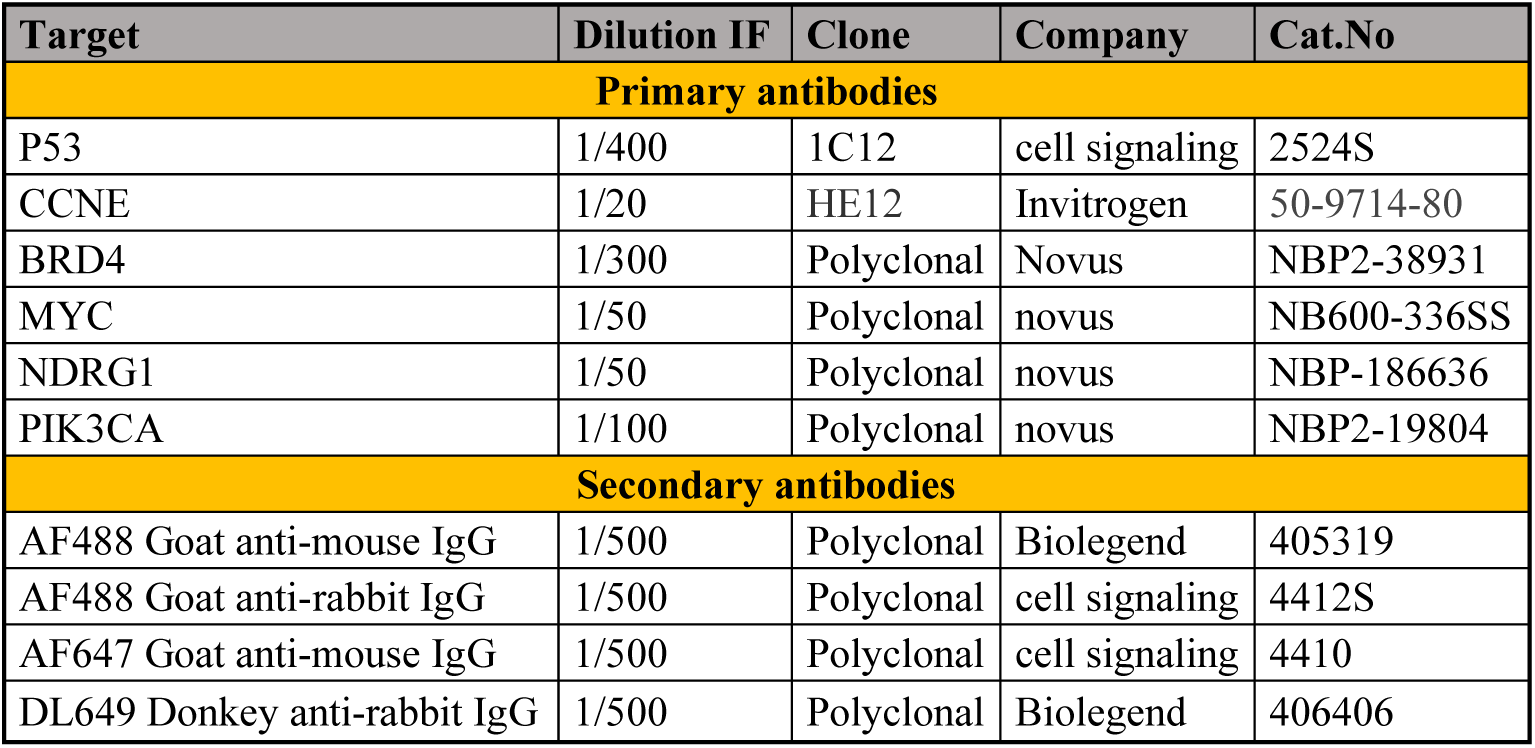
Immunofluorescence antibody panel.

**Supplementary Table 2.**
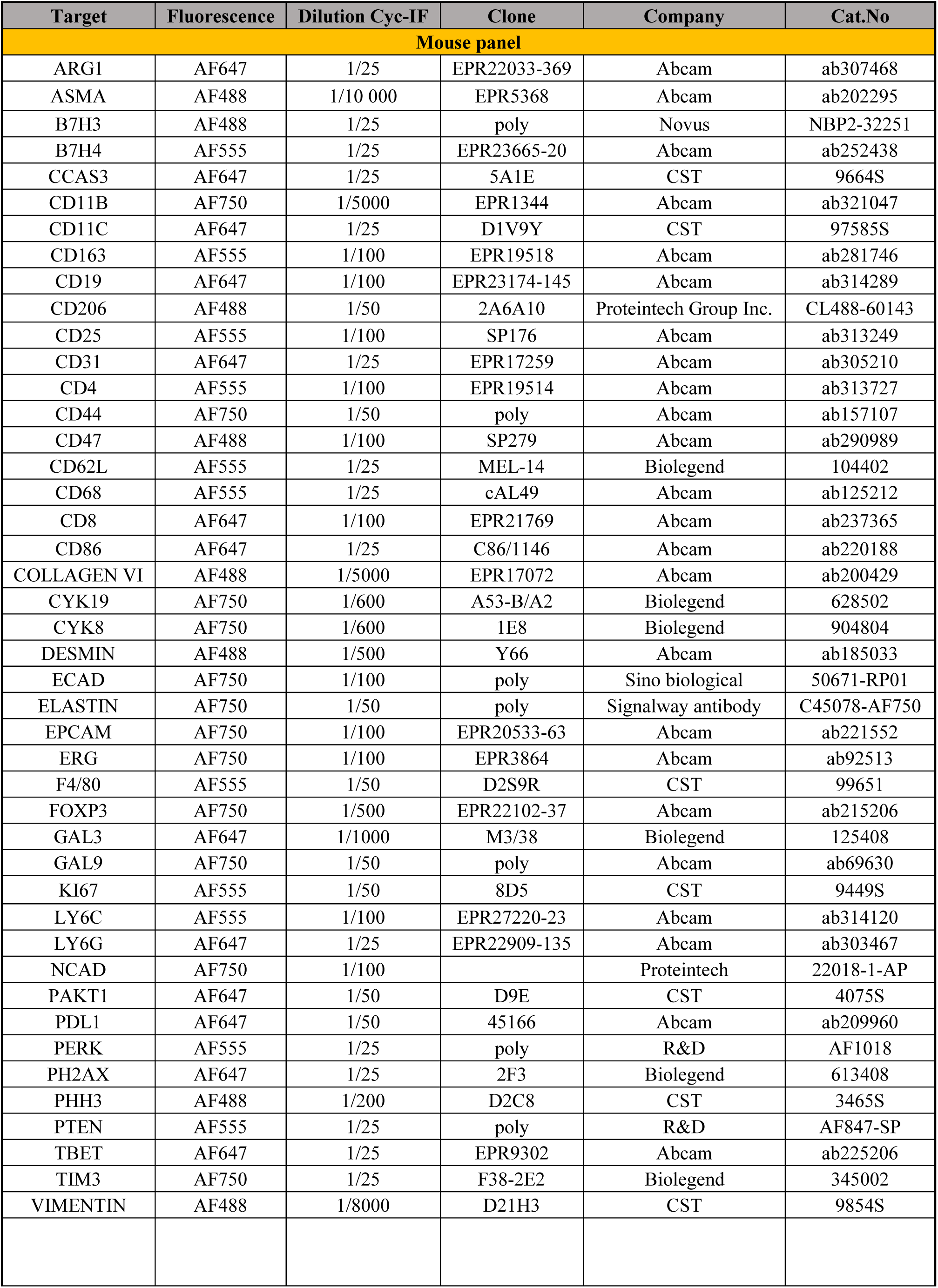
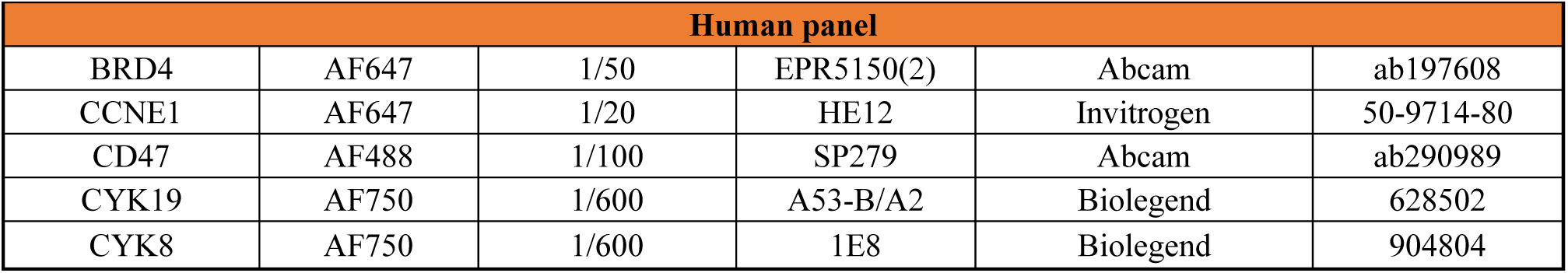
Cyc-IF antibody panel.

| Target | Fluorescence | Dilution Cyc-IF | Clone | Company | Cat.No |
| --- | --- | --- | --- | --- | --- |
| **Mouse panel** |  |  |  |  |  |
| ARG1 | AF647 | 1/25 | EPR22033-369 | Abcam | ab307468 |
| ASMA | AF488 | 1/10 000 | EPR5368 | Abcam | ab202295 |
| B7H3 | AF488 | 1/25 | poly | Novus | NBP2-32251 |
| B7H4 | AF555 | 1/25 | EPR23665-20 | Abcam | ab252438 |
| CCAS3 | AF647 | 1/25 | 5A1E | CST | 9664S |
| CD11B | AF750 | 1/5000 | EPR1344 | Abcam | ab321047 |
| CD11C | AF647 | 1/25 | D1V9Y | CST | 97585S |
| CD163 | AF555 | 1/100 | EPR19518 | Abcam | ab281746 |
| CD19 | AF647 | 1/100 | EPR23174-145 | Abcam | ab314289 |
| CD206 | AF488 | 1/50 | 2A6A10 | Proteintech Group Inc. | CL488-60143 |
| CD25 | AF555 | 1/100 | SP176 | Abcam | ab313249 |
| CD31 | AF647 | 1/25 | EPR17259 | Abcam | ab305210 |
| CD4 | AF555 | 1/100 | EPR19514 | Abcam | ab313727 |
| CD44 | AF750 | 1/50 | poly | Abcam | ab157107 |
| CD47 | AF488 | 1/100 | SP279 | Abcam | ab290989 |
| CD62L | AF555 | 1/25 | MEL-14 | Biolegend | 104402 |
| CD68 | AF555 | 1/25 | cAL49 | Abcam | ab125212 |
| CD8 | AF647 | 1/100 | EPR21769 | Abcam | ab237365 |
| CD86 | AF647 | 1/25 | C86/1146 | Abcam | ab220188 |
| COLLAGEN VI | AF488 | 1/5000 | EPR17072 | Abcam | ab200429 |
| CYK19 | AF750 | 1/600 | A53-B/A2 | Biolegend | 628502 |
| CYK8 | AF750 | 1/600 | 1E8 | Biolegend | 904804 |
| DESMIN | AF488 | 1/500 | Y66 | Abcam | ab185033 |
| ECAD | AF750 | 1/100 | poly | Sino biological | 50671-RP01 |
| ELASTIN | AF750 | 1/50 | poly | Signalway antibody | C45078-AF750 |
| EPCAM | AF750 | 1/100 | EPR20533-63 | Abcam | ab221552 |
| ERG | AF750 | 1/100 | EPR3864 | Abcam | ab92513 |
| F4/80 | AF555 | 1/50 | D2S9R | CST | 99651 |
| FOXP3 | AF750 | 1/500 | EPR22102-37 | Abcam | ab215206 |
| GAL3 | AF647 | 1/1000 | M3/38 | Biolegend | 125408 |
| GAL9 | AF750 | 1/50 | poly | Abcam | ab69630 |
| KI67 | AF555 | 1/50 | 8D5 | CST | 9449S |
| LY6C | AF555 | 1/100 | EPR27220-23 | Abcam | ab314120 |
| LY6G | AF647 | 1/25 | EPR22909-135 | Abcam | ab303467 |
| NCAD | AF750 | 1/100 |  | Proteintech | 22018-1-AP |
| PAKT1 | AF647 | 1/50 | D9E | CST | 4075S |
| PDL1 | AF647 | 1/50 | 45166 | Abcam | ab209960 |
| PERK | AF555 | 1/25 | poly | R&D | AF1018 |
| PH2AX | AF647 | 1/25 | 2F3 | Biolegend | 613408 |
| PHH3 | AF488 | 1/200 | D2C8 | CST | 3465S |
| PTEN | AF555 | 1/25 | poly | R&D | AF847-SP |
| TBET | AF647 | 1/25 | EPR9302 | Abcam | ab225206 |
| TIM3 | AF750 | 1/25 | F38-2E2 | Biolegend | 345002 |
| VIMENTIN | AF488 | 1/8000 | D21H3 | CST | 9854S |

| Human panel |  |  |  |  |  |
| --- | --- | --- | --- | --- | --- |
| BRD4 | AF647 | 1/50 | EPR5150(2) | Abcam | ab197608 |
| CCNE1 | AF647 | 1/20 | HE12 | Invitrogen | 50-9714-80 |
| CD47 | AF488 | 1/100 | SP279 | Abcam | ab290989 |
| CYK19 | AF750 | 1/600 | A53-B/A2 | Biolegend | 628502 |
| CYK8 | AF750 | 1/600 | 1E8 | Biolegend | 904804 |

**Supplementary Table 3.**
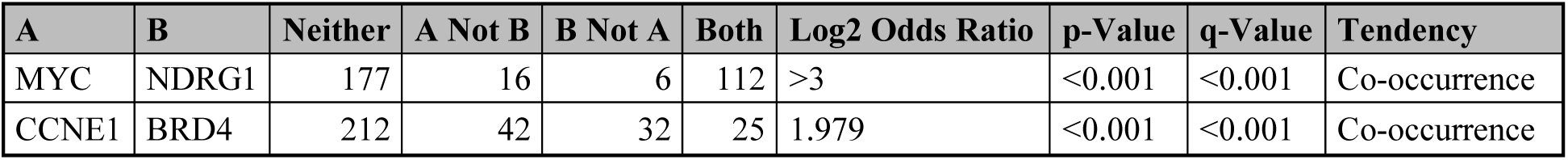
Co-occurrence between common HGSC-associated genetic alterations.

| A | B | Neither | A Not B | B Not A | Both | Log2 Odds Ratio | p-Value | q-Value | Tendency |
| --- | --- | --- | --- | --- | --- | --- | --- | --- | --- |
| MYC | NDRG1 | 177 | 16 | 6 | 112 | >3 | <0.001 | <0.001 | Co-occurrence |
| CCNE1 | BRD4 | 212 | 42 | 32 | 25 | 1.979 | <0.001 | <0.001 | Co-occurrence |

**Supplementary Table 4.** Histology features across HGSC syngeneic models.

| Type | Histology (OMS-5-2020) | Architecture |
| --- | --- | --- |
| PMN | HGSC dedifferentiated | solid, cribriform, tubular, papillary, micropapillary |
| PMN | HGSC | solid, cribriform, papillary, micropapillary |
| PMN | HGSC dedifferentiated | solid, cribriform, papillary |
| PMN | HGSC dedifferentiated | solid, cribriform, papillary |
| PC | Carcinoma undifferentiated | solid |
| PC | HGSC | solid, tubular, papillary, cribriform |
| PC | HGSC | papillary, micropapillary, tubular, cribriform |
| PCBr | HGSC dedifferentiated | solid, papillary, micropapillary, tubular |
| PCBr | HGSC dedifferentiated | solid, papillary, micropapillary, tubular |
| PCBr | HGSC dedifferentiated | solid, papillary, micropapillary, tubular |
| PCBr | HGSC dedifferentiated | solid, papillary, micropapillary, tubular |
| PBr | HGSC | solid, cribriform, tubular |
| PBr | HGSC | papillary, micropapillary, tubular, cribriform |
| PBr | HGSC | solid, cribriform, tubular, micropapillary |
| PBr | HGSC | solid, cribriform, tubular, papillary |
| PBBr | HGSC dedifferentiated | solid, tubular |
| PBBr | HGSC dedifferentiated | solid, tubular |
| PBBr | HGSC dedifferentiated | solid, tubular |
| PBBr | Carcinoma undifferentiated | solid |
| PPNM | HGSC dedifferentiated | solid, papillary |
| PPNM | HGSC dedifferentiated | solid, papillary, cribriform, tubular |
| PPNM | HGSC dedifferentiated | solid, papillary, cribriform, tubular |
| PPNM | HGSC dedifferentiated | solid, papillary, tubular |
| BPPNM | HGSC dedifferentiated | solid, tubular |
| BPPNM | Carcinoma undifferentiated | solid |
| BPPNM | HGSC dedifferentiated | solid, tubular |
| BPPNM | HGSC dedifferentiated | solid, tubular, micropapillary |
| PBMN | Carcinoma undifferentiated | solid |
| PBMN | Carcinoma undifferentiated | solid |
| PBMN | Carcinoma undifferentiated | solid |
| PBMN | Carcinoma undifferentiated | solid |

## References

1 Azzalini, E., Stanta, G., Canzonieri, V. & Bonin, S. Overview of Tumor Heterogeneity in High-Grade Serous Ovarian Cancers. Int J Mol Sci 24 (2023). 10.3390/ijms242015077

2 Cancer Genome Atlas Research, N. Integrated genomic analyses of ovarian carcinoma. Nature 474, 609–615 (2011). 10.1038/nature10166

3 Scott, C. L. et al. Ovarian cancer. Nat Rev Dis Primers 12 (2026). 10.1038/s41572-026-00686-x

4 Polajzer, S. & Cerne, K. Precision Medicine in High-Grade Serous Ovarian Cancer: Targeted Therapies and the Challenge of Chemoresistance. Int J Mol Sci 26 (2025). 10.3390/ijms26062545

5 Miller, R. E., Elyashiv, O., El-Shakankery, K. H. & Ledermann, J. A. Ovarian Cancer Therapy: Homologous Recombination Deficiency as a Predictive Biomarker of Response to PARP Inhibitors. Onco Targets Ther 15, 1105–1117 (2022). 10.2147/OTT.S272199

6 Bryant, H. E. et al. Specific killing of BRCA2-deficient tumours with inhibitors of poly(ADP-ribose) polymerase. Nature 434, 913–917 (2005). 10.1038/nature03443

7 Farmer, H. et al. Targeting the DNA repair defect in BRCA mutant cells as a therapeutic strategy. Nature 434, 917–921 (2005). 10.1038/nature03445

8 Loverix, L. et al. PARP inhibitor predictive value of the Leuven HRD test compared with Myriad MyChoice CDx PLUS HRD on 468 ovarian cancer patients from the PAOLA-1/ENGOT-ov25 trial. Eur J Cancer 188, 131–139 (2023). 10.1016/j.ejca.2023.04.020

9 Bowtell, D. D. et al. Rethinking ovarian cancer II: reducing mortality from high-grade serous ovarian cancer. Nat Rev Cancer 15, 668–679 (2015). 10.1038/nrc4019

10 da Costa, A., Chowdhury, D., Shapiro, G. I., D’Andrea, A. D. & Konstantinopoulos, P. A. Targeting replication stress in cancer therapy. Nat Rev Drug Discov 22, 38–58 (2023). 10.1038/s41573-022-00558-5

11 Tang, Q., Wang, X., Wang, H., Zhong, L. & Zou, D. Advances in ATM, ATR, WEE1, and CHK1/2 inhibitors in the treatment of PARP inhibitor-resistant ovarian cancer. Cancer Biol Med 20, 915–921 (2024). 10.20892/j.issn.2095-3941.2023.0260

12 Sun, C., Fang, Y., Labrie, M., Li, X. & Mills, G. B. Systems approach to rational combination therapy: PARP inhibitors. Biochem Soc Trans 48, 1101–1108 (2020). 10.1042/BST20191092

13 Fang, Y. et al. Sequential Therapy with PARP and WEE1 Inhibitors Minimizes Toxicity while Maintaining Efficacy. Cancer Cell 35, 851–867 e857 (2019). 10.1016/j.ccell.2019.05.001

14 Drew, Y., Zenke, F. T. & Curtin, N. J. DNA damage response inhibitors in cancer therapy: lessons from the past, current status and future implications. Nat Rev Drug Discov 24, 19–39 (2025). 10.1038/s41573-024-01060-w

15 Gupta, R., Kumar, R., Penn, C. A. & Wajapeyee, N. Immune evasion in ovarian cancer: implications for immunotherapy and emerging treatments. Trends Immunol 46, 166–181 (2025). 10.1016/j.it.2024.12.006

16 Kandalaft, L. E., Dangaj Laniti, D. & Coukos, G. Immunobiology of high-grade serous ovarian cancer: lessons for clinical translation. Nat Rev Cancer 22, 640–656 (2022). 10.1038/s41568-022-00503-z

17 Mateiou, C. et al. Spatial tumor immune microenvironment phenotypes in ovarian cancer. NPJ Precis Oncol 8, 148 (2024). 10.1038/s41698-024-00640-8

18 Sun, J. et al. Immuno-genomic characterisation of high-grade serous ovarian cancer reveals immune evasion mechanisms and identifies an immunological subtype with a favourable prognosis and improved therapeutic efficacy. Br J Cancer 126, 1570–1580 (2022). 10.1038/s41416-021-01692-4

19 Bai, Y., Chen, R. & Lu, X. A CAF-Associated Stromal Remodeling Signature Links Immune Exclusion to Exhaustion-Prone CD8(+) T-Cell Dysfunction in High-Grade Serous Ovarian Cancer. Int J Mol Sci 27 (2026). 10.3390/ijms27136092

20 Yoon, W. H., DeFazio, A. & Kasherman, L. Immune checkpoint inhibitors in ovarian cancer: where do we go from here? Cancer Drug Resist 6, 358–377 (2023). 10.20517/cdr.2023.13

21 Iyer, S. et al. Genetically Defined Syngeneic Mouse Models of Ovarian Cancer as Tools for the Discovery of Combination Immunotherapy. Cancer Discov 11, 384–407 (2021). 10.1158/2159-8290.CD-20-0818

22 Zhang, S. et al. Genetically Defined, Syngeneic Organoid Platform for Developing Combination Therapies for Ovarian Cancer. Cancer Discov 11, 362–383 (2021). 10.1158/2159-8290.CD-20-0455

23 Crespo Oliva, C. D. C., Gerber, Z., Jean, D., Allard-Chamard, H. & Labrie, M. CCR2(-) T peripheral helper cells as potential coordinators of local immune architecture in human cancer. Discov Immunol 5, kyag007 (2026). 10.1093/discim/kyag007

24 Pejovic, T. et al. Single-Cell Proteomics Analysis of Recurrent Low-Grade Serous Ovarian Carcinoma and Associated Brain Metastases. Frontiers in Oncology 12 (2022). 10.3389/fonc.2022.903806

25 Ramanathan, B. et al. Expanding the clinical spectrum of autoimmune inflammatory myopathies with prominent B cell aggregates: a case series. medRxiv, 2026.2003.2001.26347357 (2026). 10.64898/2026.03.01.26347357

26 Muhlich, J. L. et al. Stitching and registering highly multiplexed whole-slide images of tissues and tumors using ASHLAR. Bioinformatics 38, 4613–4621 (2022). 10.1093/bioinformatics/btac544

27 Gerber, Z. et al. SCORPy: Lowering the computational barrier to reproducible multiplexed imaging spatial single cell proteomics analysis. bioRxiv, 2026.2008.2024.746722 (2026). 10.64898/2026.08.24.746722

28 Etemadmoghadam, D. et al. Synthetic lethality between CCNE1 amplification and loss of BRCA1. Proc Natl Acad Sci U S A 110, 19489–19494 (2013). 10.1073/pnas.1314302110

29 Haagsma, J. et al. Gain-of-function p53(R175H) blocks apoptosis in a precursor model of ovarian high-grade serous carcinoma. Sci Rep 13, 11424 (2023). 10.1038/s41598-023-38609-5

30 Tuna, M. et al. Clinical relevance of TP53 hotspot mutations in high-grade serous ovarian cancers. Br J Cancer 122, 405–412 (2020). 10.1038/s41416-019-0654-8

31 Kotsantis, P., Petermann, E. & Boulton, S. J. Mechanisms of Oncogene-Induced Replication Stress: Jigsaw Falling into Place. Cancer Discov 8, 537–555 (2018). 10.1158/2159-8290.CD-17-1461

32 Zeman, M. K. & Cimprich, K. A. Causes and consequences of replication stress. Nat Cell Biol 16, 2–9 (2014). 10.1038/ncb2897

33 Berenjeno, I. M. et al. Oncogenic PIK3CA induces centrosome amplification and tolerance to genome doubling. Nat Commun 8, 1773 (2017). 10.1038/s41467-017-02002-4

34 Hanker, A. B., Kaklamani, V. & Arteaga, C. L. Challenges for the Clinical Development of PI3K Inhibitors: Strategies to Improve Their Impact in Solid Tumors. Cancer Discov 9, 482–491 (2019). 10.1158/2159-8290.CD-18-1175

35 Limas, J. C. et al. Quantitative profiling of adaptation to cyclin E overproduction. Life Sci Alliance 5 (2022). 10.26508/lsa.202201378

36 Takahashi, N. et al. Replication stress defines distinct molecular subtypes across cancers. Cancer Res Commun 2, 503–517 (2022). 10.1158/2767-9764.crc-22-0168

37 McCluggage, W. G., Singh, N. & Gilks, C. B. Key changes to the World Health Organization (WHO) classification of female genital tumours introduced in the 5th edition (2020). Histopathology 80, 762–778 (2022). 10.1111/his.14609

38 Denisov, E. V. et al. Clinically relevant morphological structures in breast cancer represent transcriptionally distinct tumor cell populations with varied degrees of epithelial-mesenchymal transition and CD44(+)CD24(-) stemness. Oncotarget 8, 61163–61180 (2017). 10.18632/oncotarget.18022

39 Bai, F. et al. BRCA1 suppresses epithelial-to-mesenchymal transition and stem cell dedifferentiation during mammary and tumor development. Cancer Res 74, 6161–6172 (2014). 10.1158/0008-5472.CAN-14-1119

40 Qi, Y. et al. PTEN suppresses epithelial-mesenchymal transition and cancer stem cell activity by downregulating Abi1. Sci Rep 10, 12685 (2020). 10.1038/s41598-020-69698-1

41 Zhang, L. et al. Effects of Kras activation and Pten deletion alone or in combination on MUC1 biology and epithelial-to-mesenchymal transition in ovarian cancer. Oncogene 35, 5010–5020 (2016). 10.1038/onc.2016.53

42 Karlsson, V. et al. Elevated Galectin-3 levels in the tumor microenvironment of ovarian cancer - implication of ROS mediated suppression of NK cell antitumor response via tumor-associated neutrophils. Front Immunol 15, 1506236 (2024). 10.3389/fimmu.2024.1506236

43 Li, Y. et al. Overexpression of CD47 predicts poor prognosis and promotes cancer cell invasion in high-grade serous ovarian carcinoma. Am J Transl Res 9, 2901–2910 (2017).

44 Petersen, S. et al. CCNE1 and BRD4 co-amplification in high-grade serous ovarian cancer is associated with poor clinical outcomes. Gynecol Oncol 157, 405–410 (2020). 10.1016/j.ygyno.2020.01.038

45 Gupta, N., Huang, T. T., Horibata, S. & Lee, J. M. Cell cycle checkpoints and beyond: Exploiting the ATR/CHK1/WEE1 pathway for the treatment of PARP inhibitor-resistant cancer. Pharmacol Res 178, 106162 (2022). 10.1016/j.phrs.2022.106162

46 Reddy, T. P. & Yap, T. A. Unlocking the therapeutic potential of ATR inhibitors: Advances, challenges, and opportunities in cancer therapy. Clin Transl Med 15, e70397 (2025). 10.1002/ctm2.70397

47 Liu, R. et al. CD47 promotes ovarian cancer progression by inhibiting macrophage phagocytosis. Oncotarget 8, 39021–39032 (2017). 10.18632/oncotarget.16547

48 Hardaker, E. L. et al. The ATR inhibitor ceralasertib potentiates cancer checkpoint immunotherapy by regulating the tumor microenvironment. Nat Commun 15, 1700 (2024). 10.1038/s41467-024-45996-4

49 Zhu, X. et al. Efficacy of the ATR Inhibitor Ceralasertib in Patients with ARID1A-Deficient Gynecologic and Other Solid Tumor Malignancies. Clin Cancer Res 32, 1632–1640 (2026). 10.1158/1078-0432.CCR-25-4043

50 Chen, J. & Zou, L. Replication stress in cancer: origins, consequences and therapeutic opportunities. Nat Rev Cancer (2026). 10.1038/s41568-026-00979-z

51 Curti, L. & Campaner, S. MYC-Induced Replicative Stress: A Double-Edged Sword for Cancer Development and Treatment. Int J Mol Sci 22 (2021). 10.3390/ijms22126168

52 Forment, J. V. & O’Connor, M. J. Targeting the replication stress response in cancer. Pharmacol Ther 188, 155–167 (2018). 10.1016/j.pharmthera.2018.03.005

53 Classen, S. et al. Partial Reduction in BRCA1 Gene Dose Modulates DNA Replication Stress Level and Thereby Contributes to Sensitivity or Resistance. Int J Mol Sci 23 (2022). 10.3390/ijms232113363

54 Lam, F. C. et al. BRD4 prevents the accumulation of R-loops and protects against transcription-replication collision events and DNA damage. Nat Commun 11, 4083 (2020). 10.1038/s41467-020-17503-y

55 Wang, G. et al. PTEN regulates RPA1 and protects DNA replication forks. Cell Res 25, 1189–1204 (2015). 10.1038/cr.2015.115

56 Khamidullina, A. I., Abramenko, Y. E., Bruter, A. V. & Tatarskiy, V. V. Key Proteins of Replication Stress Response and Cell Cycle Control as Cancer Therapy Targets. Int J Mol Sci 25 (2024). 10.3390/ijms25021263

57 Venegas, L. & Lheureux, S. Interplay of replication stress response and immune microenvironment in high-grade serous ovarian cancer. Front Cell Dev Biol 13, 1638964 (2025). 10.3389/fcell.2025.1638964

58 Jones, R. M. et al. Increased replication initiation and conflicts with transcription underlie Cyclin E-induced replication stress. Oncogene 32, 3744–3753 (2013). 10.1038/onc.2012.387

59 Martinikova, A. S. et al. PPM1D activity promotes the replication stress caused by cyclin E1 overexpression. Mol Oncol 18, 6–20 (2024). 10.1002/1878-0261.13433

60 Dominguez-Sola, D. & Gautier, J. MYC and the control of DNA replication. Cold Spring Harb Perspect Med 4 (2014). 10.1101/cshperspect.a014423

61 Macheret, M. & Halazonetis, T. D. Intragenic origins due to short G1 phases underlie oncogene-induced DNA replication stress. Nature 555, 112–116 (2018). 10.1038/nature25507

62 He, J., Kang, X., Yin, Y., Chao, K. S. & Shen, W. H. PTEN regulates DNA replication progression and stalled fork recovery. Nat Commun 6, 7620 (2015). 10.1038/ncomms8620

63 Hou, S. Q., Ouyang, M., Brandmaier, A., Hao, H. & Shen, W. H. PTEN in the maintenance of genome integrity: From DNA replication to chromosome segregation. Bioessays 39 (2017). 10.1002/bies.201700082

64 Huen, M. S., Sy, S. M. & Chen, J. BRCA1 and its toolbox for the maintenance of genome integrity. Nat Rev Mol Cell Biol 11, 138–148 (2010). 10.1038/nrm2831

65 Tarsounas, M. & Sung, P. The antitumorigenic roles of BRCA1-BARD1 in DNA repair and replication. Nat Rev Mol Cell Biol 21, 284–299 (2020). 10.1038/s41580-020-0218-z

66 Konstantinopoulos, P. A., Ceccaldi, R., Shapiro, G. I. & D’Andrea, A. D. Homologous Recombination Deficiency: Exploiting the Fundamental Vulnerability of Ovarian Cancer. Cancer Discov 5, 1137–1154 (2015). 10.1158/2159-8290.CD-15-0714

67 Zhao, W., Wiese, C., Kwon, Y., Hromas, R. & Sung, P. The BRCA Tumor Suppressor Network in Chromosome Damage Repair by Homologous Recombination. Annu Rev Biochem 88, 221–245 (2019). 10.1146/annurev-biochem-013118-111058

68 Sansregret, L., Vanhaesebroeck, B. & Swanton, C. Determinants and clinical implications of chromosomal instability in cancer. Nat Rev Clin Oncol 15, 139–150 (2018). 10.1038/nrclinonc.2017.198

69 Du, W., Xia, X., Hu, F. & Yu, J. Extracellular matrix remodeling in the tumor immunity. Front Immunol 14, 1340634 (2023). 10.3389/fimmu.2023.1340634

70 Vyas, M. & Demehri, S. The extracellular matrix and immunity: breaking the old barrier in cancer. Trends Immunol 43, 423–425 (2022). 10.1016/j.it.2022.04.004

71 Nakayama, N. et al. Gene amplification CCNE1 is related to poor survival and potential therapeutic target in ovarian cancer. Cancer 116, 2621–2634 (2010). 10.1002/cncr.24987

72 Gorski, J. W., Ueland, F. R. & Kolesar, J. M. CCNE1 Amplification as a Predictive Biomarker of Chemotherapy Resistance in Epithelial Ovarian Cancer. Diagnostics (Basel) 10 (2020). 10.3390/diagnostics10050279

73 Lheureux, S. et al. EVOLVE: A Multicenter Open-Label Single-Arm Clinical and Translational Phase II Trial of Cediranib Plus Olaparib for Ovarian Cancer after PARP Inhibition Progression. Clin Cancer Res 26, 4206–4215 (2020). 10.1158/1078-0432.CCR-19-4121

74 Xu, H. et al. CHK1 inhibitor SRA737 is active in PARP inhibitor resistant and CCNE1 amplified ovarian cancer. iScience 27, 109978 (2024). 10.1016/j.isci.2024.109978

75 Beers, S. A., Glennie, M. J. & White, A. L. Influence of immunoglobulin isotype on therapeutic antibody function. Blood 127, 1097–1101 (2016). 10.1182/blood-2015-09-625343

76 Vukovic, N. et al. Mouse IgG2a Isotype Therapeutic Antibodies Elicit Superior Tumor Growth Control Compared with mIgG1 or mIgE. Cancer Res Commun 3, 109–118 (2023). 10.1158/2767-9764.CRC-22-0356

77 Bastiaenen, V. P. et al. A mouse model for peritoneal metastases of colorectal origin recapitulates patient heterogeneity. Lab Invest 100, 1465–1474 (2020). 10.1038/s41374-020-0448-x

